# Conserved cardiolipin-mitochondrial ADP/ATP carrier interactions assume distinct structural and functional roles that are clinically relevant

**DOI:** 10.1101/2023.05.05.539595

**Authors:** Nanami Senoo, Dinesh K. Chinthapalli, Matthew G. Baile, Vinaya K. Golla, Bodhisattwa Saha, Oluwaseun B. Ogunbona, James A. Saba, Teona Munteanu, Yllka Valdez, Kevin Whited, Dror Chorev, Nathan N. Alder, Eric R. May, Carol V. Robinson, Steven M. Claypool

## Abstract

The mitochondrial phospholipid cardiolipin (CL) promotes bioenergetics via oxidative phosphorylation (OXPHOS). Three tightly bound CLs are evolutionarily conserved in the ADP/ATP carrier (AAC in yeast; adenine nucleotide translocator, ANT in mammals) which resides in the inner mitochondrial membrane and exchanges ADP and ATP to enable OXPHOS. Here, we investigated the role of these buried CLs in the carrier using yeast Aac2 as a model. We introduced negatively charged mutations into each CL-binding site of Aac2 to disrupt the CL interactions via electrostatic repulsion. While all mutations disturbing the CL-protein interaction destabilized Aac2 monomeric structure, transport activity was impaired in a pocket-specific manner. Finally, we determined that a disease-associated missense mutation in one CL-binding site in ANT1 compromised its structure and transport activity, resulting in OXPHOS defects. Our findings highlight the conserved significance of CL in AAC/ANT structure and function, directly tied to specific lipid-protein interactions.

## Introduction

Biological membranes consist heterogeneously of proteins and lipids. Mitochondria contain two membranes, the outer mitochondrial membrane (OMM) and the inner mitochondrial membrane (IMM). While the OMM compartmentalizes the organelle by forming its outer edge, the IMM forms infoldings called cristae which concentrate the bioenergetic components required for oxidative phosphorylation (OXPHOS) ^1–4^. Cardiolipin (CL) is a phospholipid enriched in mitochondria, especially in the IMM^5^. Consisting of two phosphate headgroups and four acyl-chains, the unique structure of CL supports and optimizes many biological events occurring in mitochondria ^6^. Focused on the OXPHOS machinery, the main documented roles of CL include physically associating with individual components ^7–20^ and stabilizing their higher order assembly into supercomplexes ^21–23^. Loss of CL diminishes energetic capacity in multiple organisms ^24–28^. This implies that conserved CL-protein interactions serve critical and functionally important roles for the OXPHOS machinery. Although the significance of CL in mitochondrial biology has been increasingly accepted, mechanistic details of how this lipid supports membrane protein structure and function are still lacking.

The ADP/ATP carrier (AAC in yeast; adenine nucleotide translocator, ANT in humans) belongs to the largest solute carrier family, SLC25, which includes 53 members in humans. SLC25 family members are mostly located in the IMM where they collectively mediate the flux of a wide variety of solutes, such as ions, cofactors, and amino acids, across the otherwise impermeable IMM ^29^. AAC is embedded in the IMM and transports ATP out of and ADP into the mitochondrial matrix in a 1:1 exchange. This function is required for OXPHOS and needed to make ATP synthesized in the matrix bioavailable to the rest of the cell. Of the three isoforms in yeast, Aac2 is the most abundant and only isoform required for OXPHOS^30^. AAC alters its conformation during the transport cycle. Inhibitors are available to trap the carrier into two extreme conformational states: carboxyatractyloside (CATR) locks the carrier in the cytosolic open state (c-state) and bongkrekic acid (BKA) locks it in the matrix open state (m-state) ^7, 31^.

First demonstrated via ^31^-P NMR ^10^ and subsequently confirmed in all of the solved crystal structures of AAC locked in the c-state with CATR, including yeast Aac2 ^7^ and bovine ANT1 ^8, 9^, AAC contains three tightly bound CL molecules. In the m-state structure of AAC with BKA, one of the three CLs remains bound to AAC whereas the second and third are lost ^31^, suggesting that the affinities of the CL-binding sites are conformation-sensitive. Protein thermostability assays have shown that CL enhances the stability of yeast Aac2 ^32^. Recent molecular dynamics (MD) simulation studies indicate that CL affects the structure of AAC/ANT ^33, 34^. Our previous study further emphasized the diverse structural roles provided by CL: CL supports Aac2 monomeric structure and its transport-related conformation ^35^. In addition, through a distinct mechanism, CL regulates the association of Aac2 with other OXPHOS components ^24, 35^.

The fact that the three CLs are bound to AAC in an evolutionally conserved manner underscores the functional significance of these lipid-protein interactions; however, molecular-level insight into if and how these CL molecules contribute to AAC structure and function has not been experimentally interrogated. In this study, we engineered a series of mutations to disrupt specific CL interactions in yeast Aac2 and experimentally verified that the lipid-protein interactions are structurally important. Interestingly, while each bound CL promotes Aac2 transport, one lipid-protein interaction is functionally essential. Finally, we provide evidence that a pathologic mutation in *ANT1* tied to human disease is structurally and functionally compromised due to the disruption of one of the three conserved tightly bound CLs. To our knowledge, this is the first documented disease-associated missense mutation whose defect derives from a disturbed interaction between a mutant protein and a specific lipid.

## Results

### Engineered CL-binding Aac2 mutants

The solved crystal structures of AAC in the c-state all include three tightly bound CL molecules ^7–9^. In each CL-binding pocket, the phosphate headgroup of CL engages the backbone peptide. Aiming to disrupt the lipid-protein interaction via electrostatic repulsion, we introduced negatively charged amino acids in the immediate vicinity of each CL-binding pocket of yeast Aac2. We generated a total of 10 single mutants―two designed to disrupt the interaction with CDL801 (pocket 1: I51E and G69D), three to disrupt the interaction with CDL800 (pocket 2: N90E, L155E, and G172E), and four to disrupt the interaction with CDL802 (pocket 3: L194E, R191D, M255E, and G267E)―and one double mutant (Fig. 1A). Every mutant was expressed at an equal level to WT Aac2 as re-introduced into the *aac2*Δ parental cells (Fig. 1B). While every mutant was able to grow on respiratory media (YPEG) at 30°C, in which cellular energy production relies on OXPHOS, the respiratory growth capacity varied (Fig. 1C). The respiratory growth capacity of *crd1*Δ and some of the Aac2 mutants was exacerbated at elevated temperature (37°C) (Fig. 1C), suggesting the involvement of CL and potentially CL-Aac2 in yeast adaptation to stress.

**Fig. 1:**
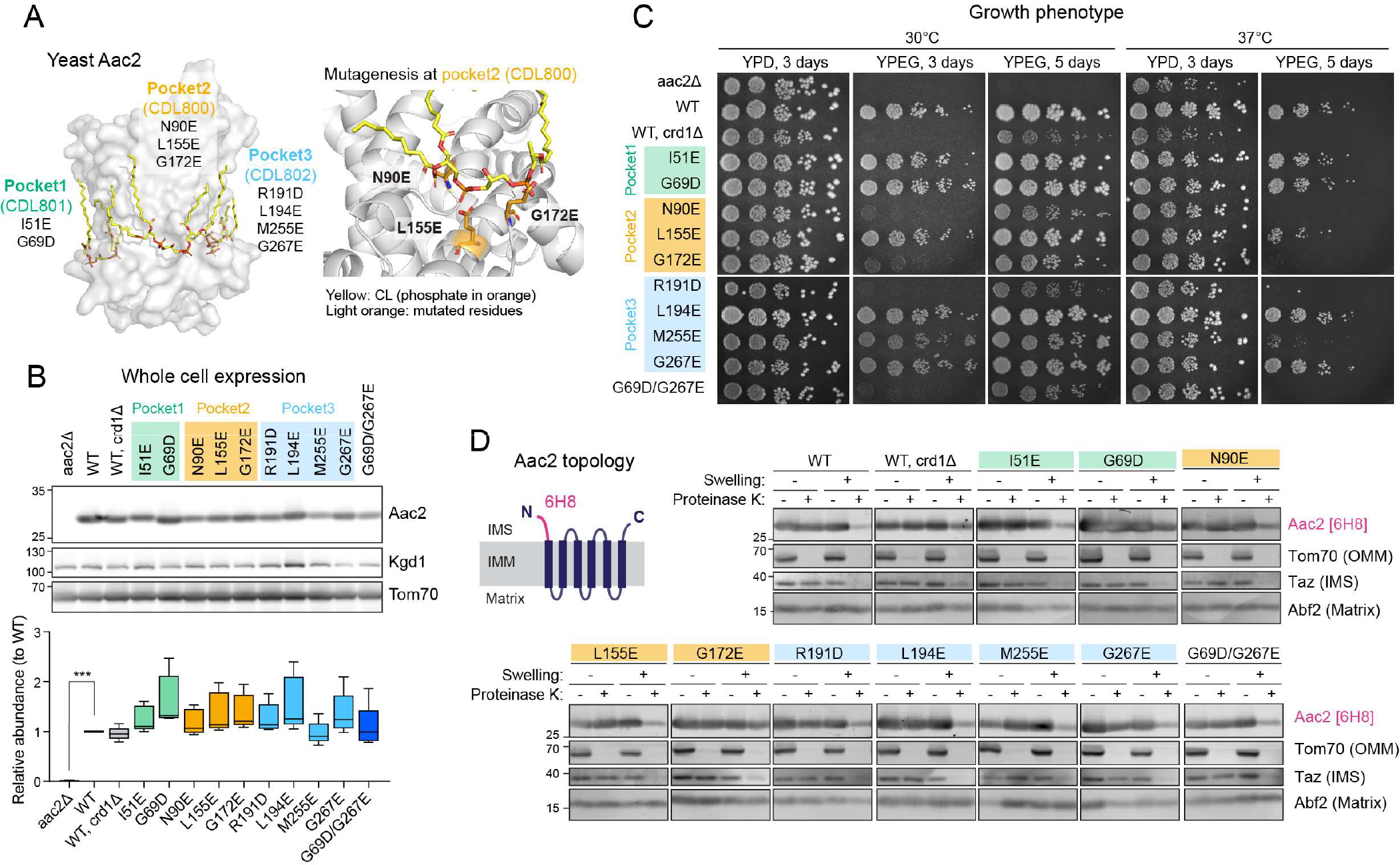
Generation and characterization of yeast Aac2 CL-binding mutants. (A) Strategy to disrupt CL-binding sites in Aac2. Aac2 was modeled onto bovine ANT1 (PDB ID: 2C3E) using SWISS-MODEL. Negatively charged amino acids introduced (light orange) into Aac2 CL-binding motifs face toward the CL phosphate headgroup (orange). (B) Expression of WT and mutant Aac2 was detected in whole cell extracts by immunoblot; Kgd1 and Tom70 served as loading controls (n=5). Significant difference was obtained by one-way ANOVA with Dunnett’s multiple comparisons test (vs. WT) ***p<0.001. (C) Growth phenotype of Aac2 CL-binding mutants. Serial dilutions of indicated cells were spotted onto fermentable (YPD) and respiratory (YPEG) media and incubated at 30 or 37°C for 3-5 days (n=3). (D) Membrane topology of WT and mutant Aac2. Isolated mitochondria were osmotically ruptured and treated with or without proteinase K as indicated. Aac2 N-terminus was detected by a monoclonal antibody 6H8 recognizing the first 13 amino acids MSSNAQVKTPLPP (n=6). IMS, intermembrane space; IMM, inner mitochondrial membrane; OMM, outer mitochondrial membrane. In B-D, representative images from the indicated replicates are shown.

Aac2 contains six membrane-passing domains with the N- and C- termini exposed to the intermembrane space ^9^. When the outer membrane was removed by swelling, proteinase K treatment erased the IMS-exposed N-terminus of Aac2 which was detected by a monoclonal antibody recognizing this peptide ^36^. With the possible exception of G172E, each Aac2 mutant presented the same pattern as WT Aac2 (Fig. 1D). This data demonstrates that the inserted negatively charged amino acids do not disrupt the topology of the mutant Aac2 proteins in the IMM.

### Specific lipid-protein interactions are disrupted in CL-binding Aac2 mutants

To identify the lipid-protein interactions of Aac2, we established a native mass spectrometry (MS) approach as our prior study successfully detected the interactions between bovine ANT1 and CLs ^37^. We introduced a Flag-tag onto the N-terminus of WT and mutant Aac2 and purified FlagAac2 from mitochondria using the detergent undecylmaltoside (UDM). Before MS analysis, the purified protein was rapidly buffer-exchanged into lauryldimethylamine oxide (LDAO). The growth phenotype of the Flag-tagged mutants was the same as untagged mutants (fig. S1A, Fig. 1C), indicating the Flag-tag is functionally silent. We noted that the use of UDM during purification was structurally destabilizing to WT Aac2; on blue-native PAGE, a smear above ∼200 kDa was seen (fig. S1B) instead of stabilized Aac2 monomer at ∼140 kDa^35^. Given this, we treated mitochondria with CATR before UDM-solubilization which indeed preserved the major ∼140 kDa Aac2 monomer, similar to what is observed following digitonin extraction (fig. S1B). Consistently, CATR pre-treatment improved the purification efficiency of FlagAac2.

On native MS analysis, WT Aac2 was detected as multiple interaction forms in the membrane that contained CL (Fig. 2A). A typical native mass spectrum of WT Aac2 generated ions in the mass range *m/z* 3500-6000 with 8+ as an abundant charge state. The mass spectrum consisted of well-resolved ions corresponding to apo Aac2, CATR bound and CATR-CL bound peaks along with very small proportions of CL bound Aac2 peaks (Fig. 2A). A maximum of CATR-bound Aac2 + 3 CLs peaks was present with a mass difference of ⁓2180Da, ⁓3580Da, and ⁓4950Da respectively. Manually conducting further analysis of the 8+ charge state, peaks that were assigned to apo Aac2 and CATR-bound Aac2 had narrow mass distribution over an *m/z* of ⁓3Da and ⁓4Da respectively, indicating a single mass composition. However, peaks assigned to CL bound Aac2 showed homogenous mass distribution over the *m/z* range of ⁓16Da, ⁓18Da, and ⁓21Da. These observations indicate that the CLs are bound in the three binding pockets when the Aac2 is stabilized in the c-state, consistent with the reported crystal structures with CATR ^7–9^. Additionally, MSMS experiments were performed on CATR-bound Aac2 + 3 CLs peak; at 150V of high collision dissociation (HCD) energy, the first CL was released from the complex followed by the second and third CL’s release with a further increase of 50V (fig. S2). Sequential loss of ⁓1400Da, which corresponds to the average mass of CL, was observed in both cases (fig. S2). Upon further increase of HCD energy to 270V, CLs were completely dissociated as peaks corresponding to CATR-bound and apo Aac2 were only observed (fig. S2). The data imply that the CL binding to one of the pockets is weaker than the other two CL binding pockets. Loss of 2 CLs at similar energy of 150∼200V implies CLs binding at two other binding pockets might have relatively similar binding strength (fig. S2). The acyl chains of CL co-purified with Aac2 were heterogeneous (CL64:4, CL 66:4, CL68:4) (fig. S3) and mirrored the natural CL profile of yeast ^25, 38^. Due to this heterogeneity, we could only predict that CL68:4 contained 16:1 and 18:1 (estimated as 16:1-16:1-18:1-18:1) as the chain lengths matched with the obtained MSMS spectra (fig. S3).

**Fig. 2:**
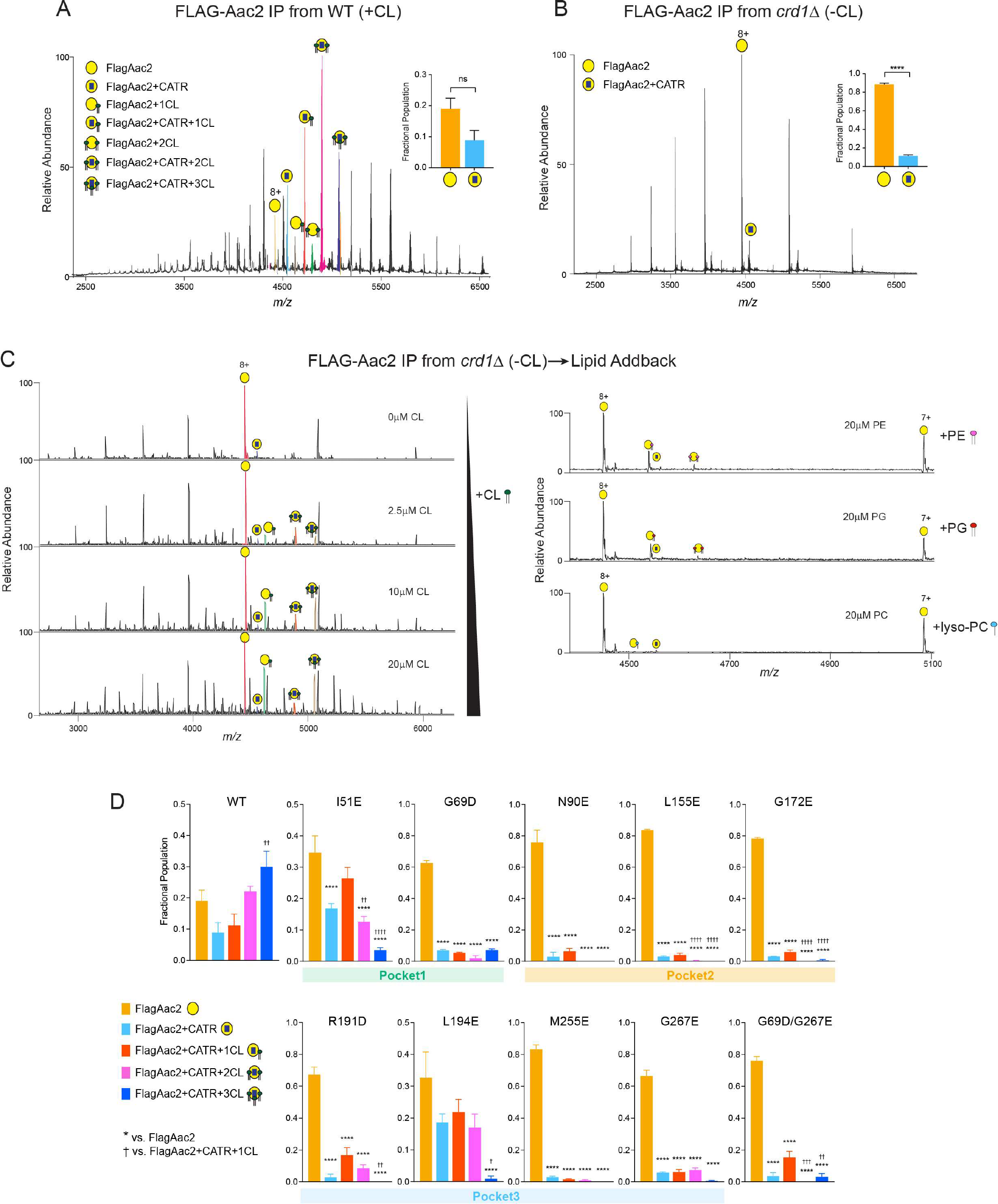
Native mass spectrometry analysis to detect lipid-protein interaction of Aac2. (A) FlagAac2 affinity purified from WT mitochondria was associated with CATR and up to three CL molecules. (B) FlagAac2 affinity purified from *crd1*Δ mitochondria which lack CL did not co-purify other phospholipids. Insets in A and B show the fractional population of FlagAac2 relative to FlagAac2+CATR when immunoprecipitated from WT and *crd1*Δ mitochondria, respectively. Mean with SEM (n=3). Significant differences as determined by Student’s t-test indicated (****p<0.0001). (C) CL, PE, PG, or lyso-PC was added to FlagAac2 purified from *crd1*Δ mitochondria at the indicated concentrations. (D) Fractional population of FlagAac2, FlagAac2+CATR alone or associated with one to three CL molecules was determined for WT and Aac2 CL-binding mutants. Mean with SEM (n=5-6). Significant differences were obtained by one-way ANOVA with Dunnett’s multiple comparisons test (* vs. FlagAac2; † vs. FlagAac2+CATR+1CL); *p<0.05, **p<0.01, ***p<0.001, ****p<0.0001.

In contrast, native MS of WT Aac2 purified from *crd1*Δ mitochondria which lack CL yielded ions corresponding to apo and CATR-bound Aac2 only; neither CL nor any other phospholipid including the CL precursor, phosphatidylglycerol (PG) which accumulates in the *crd1*Δ strain ^25^, were co-purified with Aac2 (Fig. 2B). We also examined the capacity of Aac2 in *crd1*Δ to recoup CL binding by adding CL exogenously. The interactions of added CL to Aac2 occurred in a titrated manner: minimal CL binding was achieved when incubated with 1 μM CL, and 2-3 CLs were bound at 2.5 μM (Fig. 2C). This ability was specific to CL and not shared by other phospholipids including PG, lyso-phosphatidylcholine (lyso-PC), and phosphatidylethanolamine (PE) (Fig. 2C). These results demonstrate that Aac2 lipid binding is highly specific to CL and that other lipid classes are unable to structurally replace CL as contained in the IMM or added exogenously. Note that an increase in number of charge state distributions in *crd1*Δ mitochondria (Fig. 2B) indicated that Aac2 is denatured or partially unfolded due to the absence of CL. Given this, negatively charged phospholipids could bind at higher concentrations, however, these are likely nonspecific.

To account for the relative strengths/extent of disruption of CL-Aac2 interactions with our mutational strategy, the fractional populations of the peaks from the mass spectra were calculated from intensities (Fig. 2D; see methods). In WT Aac2, a high population of CATR- bound Aac2 + 3 CLs was observed (Fig. 2D), as expected for the CATR-bound form ^7–9^. For all mutants, the populations of apo Aac2 were the highest with a dramatic reduction in the populations of CATR-bound Aac2 + 3 CLs (Fig. 2D), demonstrating the feasibility of our mutational strategy. Though this is evident in all the mutants, the extent of disruption of CATR- bound Aac2 + 3 CLs was less in the mutants generated in pocket 1 (I51E and G69D) and double mutant (G69D/G267E). Prominent destabilization of CL-protein interactions was bestowed by the mutants of pocket 2 wherein fractional populations of CATR-bound Aac2 + 2 CLs were also significantly reduced. This disruption of CATR-bound Aac2 + 2 CLs was also seen in the double mutant (G69D/G267E) (Fig. 2D). Interactions of CATR-bound Aac2 + 1 CL were retained in all the mutants and with two exceptions (G69D and G69D/G267E) trended to be higher than the 3 CLs bound form. Though the abundances of the CATR-bound with CLs species were significantly decreased in all the mutants except I51E and L194E, none of the mutants disrupted all three CATR-stabilized Aac2-CL interactions completely. Naively, we predicted that the population of CATR-bound Aac2 + 3 CLs would be the most reduced for each mutant as the inserted mutation targeted only one CL-binding pocket; in principle, the other two pockets should be intact and CL-binding competent. Although this pattern was obtained for I51E and L194E, the majority of mutants displayed more broadly disrupted CL-binding. This could indicate that ablation of any single tightly bound CL weakens the tertiary stability of mutant Aac2 and that the application of additional structurally perturbing agents, such as the harsh detergents used for native MS sample preparation, causes mutants to denature and thus release most of the bound CLs.

### CLs directly bound to Aac2 stabilize its monomeric structure

We previously demonstrated that CL stabilizes the Aac2 monomeric folding structure ^35^ but whether this property derived from one or more of the co-crystallized CL molecules was unresolved. To ask whether the CL molecules directly interacting with Aac2 affect its folded structure, we solubilized WT and mutant mitochondria with the mild detergent digitonin and resolved them on blue native-PAGE. Consistent with our previous results ^35^, the majority of WT Aac2 formed a stable monomer (locked at ∼140 kDa) in the presence of CL (lane 2 in Fig. 3A, B), whereas in the absence of CL (*crd1*Δ), the Aac2 monomer was destabilized and detected as a smear from ∼230 to < 67 kDa (lane 4 in Fig. 3A, B). Importantly, all Aac2 mutants were destabilized even in the presence of CL (even numbered lanes starting from 6 in Fig. 3A, B); this strongly indicates that, as we hypothesized, all three bound CLs are crucial in stabilizing the Aac2 monomeric structure.

**Fig. 3:**
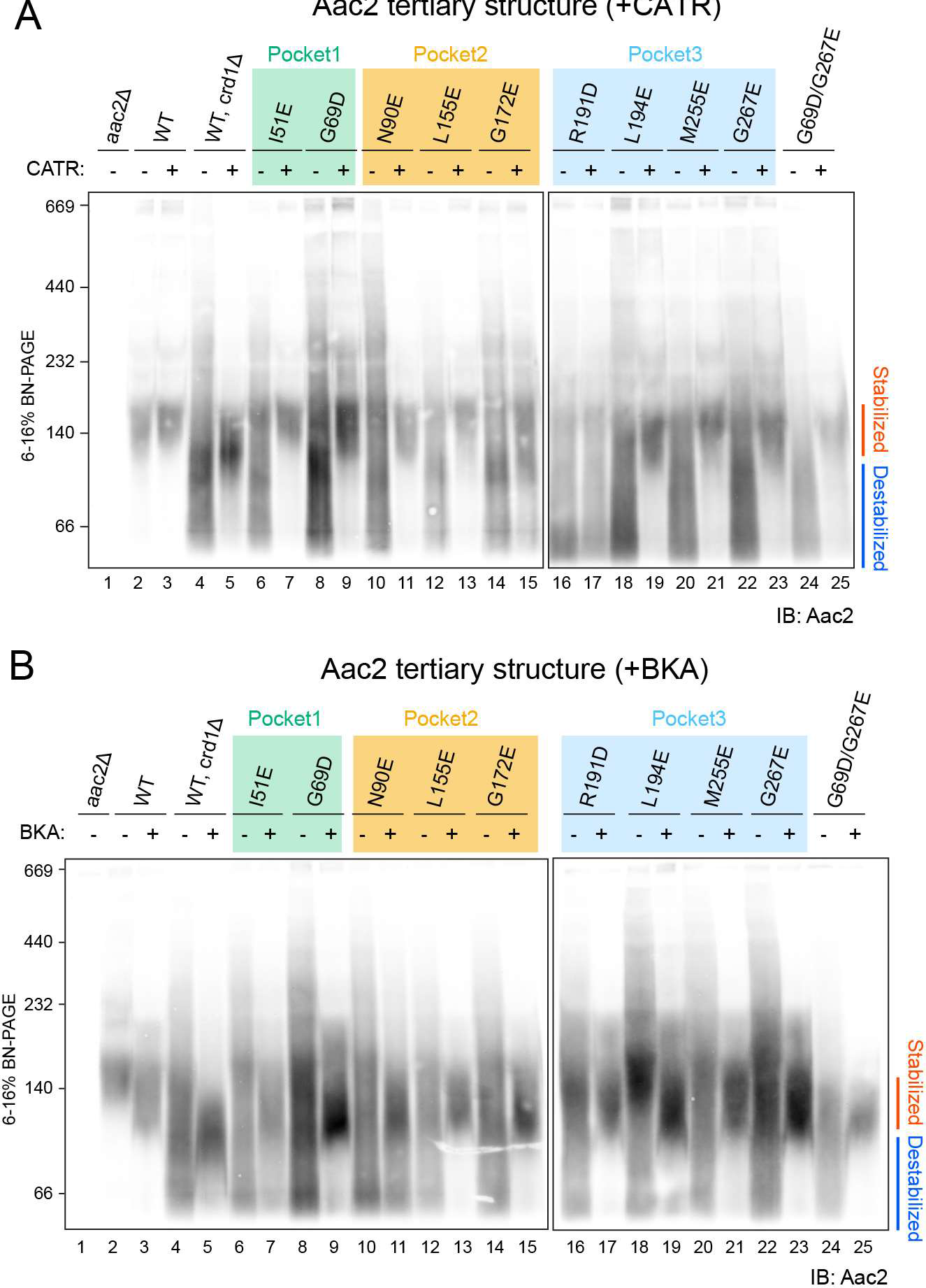
CLs associated with Aac2 stabilize its monomeric structure. Mitochondria (100 μg) from WT and Aac2 CL-binding mutants were mock-treated or instead incubated with either 40 μM CATR (**A**) or 10 μM BKA (**B**) and then solubilized with 1.5% (w/v) digitonin. The extracts were resolved by 6 to 16% blue native-PAGE and immunoblotted for Aac2 (n=6). Representative images from the indicated replicates are shown.

We also tested the effects of the inhibitors CATR and BKA, which tightly lock the carrier in distinct c-state and m-state conformations, respectively^7, 31^. As seen previously^35^, pre-treatment of *crd1*Δ mitochondria with either CATR or BKA fixed the Aac2 monomeric structure (lane 5 in Fig. 3A, B), demonstrating that Aac2 is properly folded and inserted into the IMM even in the absence of CL because CATR and BKA do not bind unfolded AAC^32^. The addition of CATR pre-digitonin solubilization rescued the predominant ∼140 kDa monomeric Aac2 c-state conformer of each CL-binding mutant (odd numbered lanes starting from 7 in Fig. 3A) save one, R191D (lane 17 in Fig. 3A). Intriguingly, the addition of BKA pre-digitonin solubilization did rescue the predominant ∼120 kDa monomeric Aac2 m-state conformer of the mutants including R191D (odd numbered lanes starting from 7 in Fig. 3B), similar to WT Aac2 in mitochondria lacking CL (lane 5 in Fig. 3B). Taken together with Fig. 1D, these results demonstrate that every CL-binding Aac2 mutant is properly folded in the IMM and suggest that interfering with the interaction of only one of the three buried CL molecules is sufficient to significantly destabilize the tertiary fold of Aac2 under blue native-PAGE conditions.

### The three tightly bound CLs are functionally distinct

To clarify if the bound CL molecules are important for Aac2 transport activity, we measured ADP/ATP exchange in isolated mitochondria from CL-binding Aac2 mutants. ADP/ATP exchange was initiated by the addition of ADP and the rate of ATP efflux was measured^39–41^. The respiratory substrates malate and pyruvate were included in the reaction to establish a robust proton motive force across the IMM which is known to stimulate AAC-mediated transport^42^. The initial velocities over a range of ADP concentrations demonstrated that, as expected, WT Aac2 had the highest capacity for ATP efflux, i.e. Aac2 transport activity (Fig. 4A, B). The mutants, except L194E, had reduced Aac2 transport activity (Fig. 4A, B). The kinetics of ATP efflux was further evaluated by Michaelis-Menten equation (Fig. 4C). Compared to WT Aac2, the mutants had higher Km values for ADP, which represents the substrate concentration at which half of the maximum reaction velocity (Vmax) is reached; this suggests that the mutants are less prone to mediate ADP/ATP exchange. There was a difference in the Aac2 transport activity across the three CL-binding pockets: the most severe defects were observed in pocket 2 mutants, followed by pocket 3 mutants with the exception of L194E. Although pocket 1 mutants retained near WT transport activity, the double pocket 1/3 mutant (G69D/G267E) showed a stronger defect in Aac2 transport activity than either corresponding single mutant. In the absence of respiratory substrates, mutant transport was much slower than when the mitochondria were energized, as expected, and the values between replicates were more variable. Despite this variability, the same trend as for energized mitochondria was still observed (fig. S4).

**Fig. 4:**
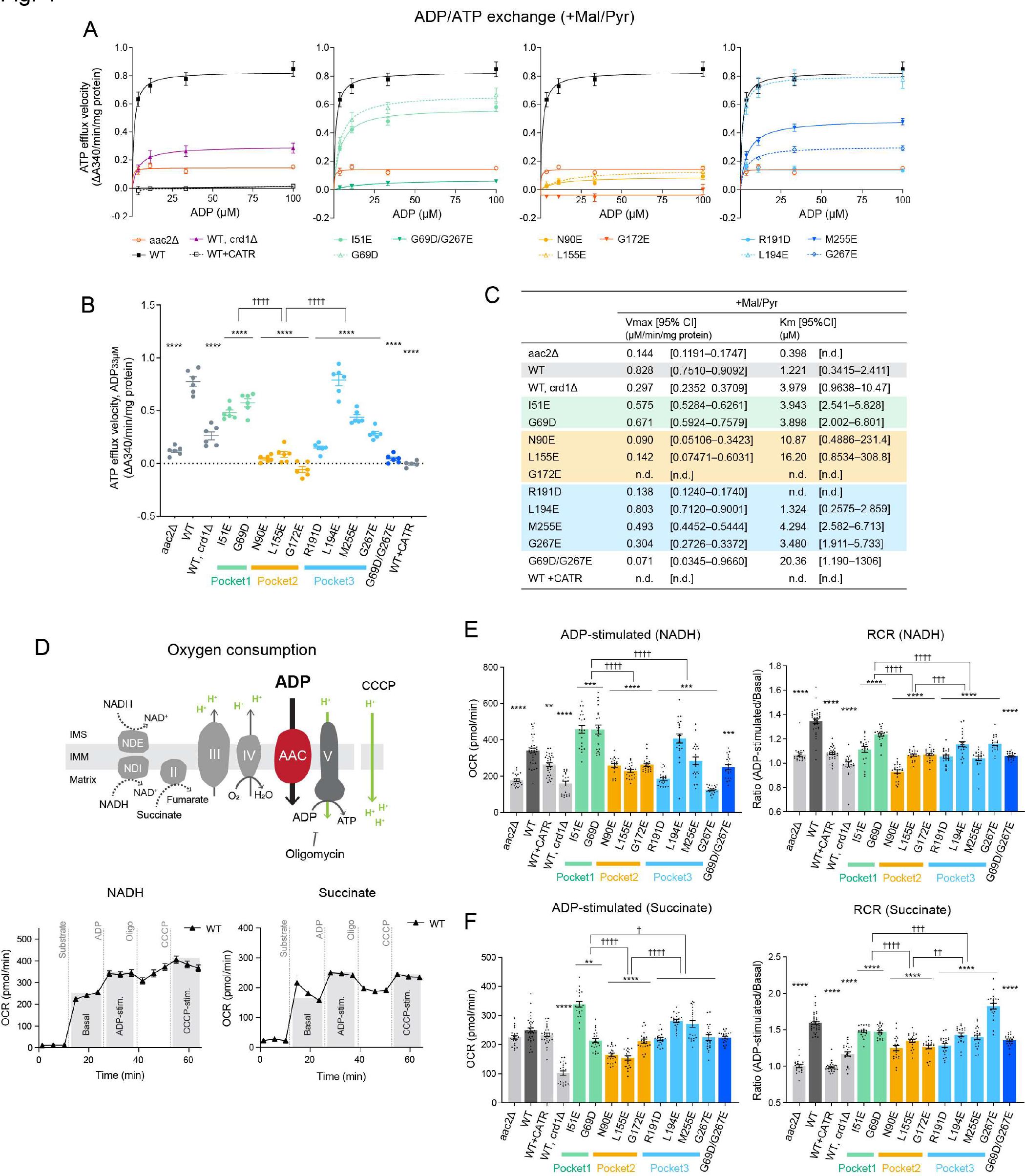
Distinct functional roles of the three tightly bound CLs. (A-C) ADP/ATP exchange: The efflux of matrix ATP was detected with isolated mitochondria as NADPH formation (A340; absorbance at 340 nm) occurring coupled with an *in vitro* glycolysis reaction which contained glucose, hexokinase, and glucose-6-phosphate dehydrogenase. The reaction was initiated by adding ADP. The measurement was performed in the presence of 5 mM malate and 5 mM pyruvate (+Mal/Pyr). Where indicated, WT mitochondria were treated with 5 μM CATR prior to the efflux reaction. (A) The linear part of the initial velocity for the ATP efflux was plotted and curve fitting performed by nonlinear regression (mean with SEM, n=6). Plots of *aac2*Δ and WT are repeated in all panels. (B) The initial velocity following the addition of 33 μM ADP presented as scatter plots (mean with SEM, n=6). (C) Fitted Km and Vmax values were obtained using the Michaelis-Menten equation (mean). The range of determined values is shown in brackets (n.d., not detected). (D) Oxygen consumption rate (OCR) in isolated mitochondria with sequential injections of respiratory substrate (NADH or succinate), ADP, oligomycin, and CCCP was measured using Seahorse XF96e FluxAnalyzer. (E, F) ADP-stimulated respiration and respiratory control ratio (RCR) of WT and Aac2 mutant mitochondria under NADH (E) and succinate (F) were plotted. RCR was obtained by dividing OCR of ADP-stimulated respiration by that of basal respiration (see also fig. S5). WT mitochondria was treated with 50 μM CATR before the measurement. Mean with SEM, n=21-35. Significant differences obtained by two-way ANOVA followed by Tukey’s multiple comparisons test are shown as * for comparison with WT and † for comparison between pocket mutants; *p<0.05, **p<0.01, ***p<0.001, ****p<0.0001.

Aac2-associated OXPHOS capacity was assessed by measuring oxygen consumption with isolated mitochondria. We sequentially injected a respiratory substrate (NADH or succinate) and ADP using a Seahorse Flux Analyzer (Fig. 4D-F, fig. S5). ADP-stimulated oxygen consumption reflects AAC transport activity as this is dependent on ADP import across the IMM through AAC. In the presence of NADH, pocket 2 and pocket 3 mutants showed reduced ADP-stimulated respiration compared to WT Aac2 (Fig. 4E). Energized with succinate, pocket 2 mutants had reduced ADP-stimulated respiration (Fig. 4F). Similar trends were observed for each respiratory substrate following the addition of the uncoupler, CCCP (fig. S5). In addition, we calculated the respiratory control ratio (RCR), defined by the ratio of oxygen consumption while ADP is phosphorylated (ADP-stimulated) to that in the absence of ADP (basal), which provides insight into OXPHOS coupling and the possibility of proton leak. As expected, in the absence of ADP/ATP exchange (*aac2*Δ and WT+CATR), RCR was ∼1 under both NADH and succinate conditions (Fig. 4E, F). With both substrates, the RCR of all pocket mutants was lower than WT; however, pocket-specific differences were detected, with RCR values decreasing for pockets 1, 3, and 2, in that order, relative to WT (Fig. 4E, F). Taken together, our data indicate that all three CL molecules bound to Aac2 regulate its transport activity but that the three buried CLs are functionally distinct: Pocket 2 has the preeminent role in supporting AAC transport activity, at least in the exchange modes tested.

### Disturbed CL-Aac2 interaction attenuates respiratory complex expression and assembly

Yeast complex IV includes three subunits encoded in the mitochondrial DNA (Cox1, Cox2, and Cox3). Using a transport-dead Aac2 mutant, we previously demonstrated that AAC transport supports complex IV expression via modulating translation of its mitochondrial DNA-encoded subunits ^43^. In this context, we noted that the mitochondrial steady-state expression of Cox1 and Cox3 were modestly reduced for those CL-binding Aac2 mutants whose transport activity was impaired the most (Fig. 4 and fig. S6A, B). This result reinforces our conclusion that the CL-Aac2 interactions are critical for AAC transport function. Even so, these mutants maintained the expression of nuclear genome-encoded respiratory complex subunits (complex III, complex IV, and complex V) as well as the mitochondrial DNA-encoded complex V subunit, Atp6.

Since the ability of Aac2 to interact with respiratory complex subunits is sensitive to its transport-related conformation which is in turn influenced by CL ^35^, we biochemically interrogated the Aac2-respiratory supercomplex (RSC) interaction for each CL-binding Aac2 mutant. As shown in fig. S7, blue native-PAGE results illustrated the higher-order protein assembly including Aac2 and RSC. In the absence of CL (*crd1*Δ), the WT Aac2-RSC interaction was significantly destabilized (fig. S7A, B), as expected ^24, 35^. While less noticeable than WT Aac2 in *crd1*Δ, the assembly of the Aac2 CL-binding mutants with RSC was compromised to varying extents (fig. S7A, B). Quantification indicated that the abundance of Aac2-III2IV2 was reduced in I51E (pocket 1), all three pocket 2 mutants, three pocket 3 mutants (R191D, M255E, G267E), and the double mutant G69D/G267E (fig. S7B). This suggests that disturbed CL-Aac2 interactions have a mild but measurable impact on Aac2-RSC interaction. While the abundance of RSCs (III2IV2 and III2IV1) was retained in all mutants (fig. S7C, D), the fraction of III2IV2 was lowered (fig. S7E). To further investigate direct interactions between Aac2 and RSC subunits, we performed Flag co-immunoprecipitation and used CATR as a folding stabilizer prior to the detergent solubilization. The pocket 2 mutant, N90E, barely interacted with the RSC subunits tested (fig. S8). Impaired protein-protein interactions were also observed for L155E and G172E (pocket 2) and R191D (pocket 3) (fig. S8C). The protein assembly defects consistently observed in the blue native-PAGE and co-immunoprecipitation results may stem from their severely weakened conformational stability and/or the relatively strong transport defect of these mutants. In fact, mutants with the most compromised transport co-purified the least amount of complex III and IV subunits (fig. S8).

### Disturbed CL-ANT1 binding underpins an uncharacterized pathogenic mutation

More than half of the CL-binding mutant residues that we targeted in yeast Aac2 are conserved in mammalian ANTs (fig. S9). Among the conserved residues, the uncharacterized L141F missense mutation in human ANT1 – corresponding to yeast Aac2 L155 in pocket 2 – was recently identified in a patient suffering from mitochondrial myopathy, exercise intolerance, hyperlactatemia, and cardiomyopathy ^44^. Since we observed that all yeast Aac2 mutants had destabilized monomeric structure (Fig. 3) and that L155 in pocket 2 was one of the crucial residues for Aac2 transport activity (Fig. 4), we hypothesized that the pathological mechanism behind this patient mutation may stem from disturbed lipid-protein interaction-based AAC/ANT dysfunction. To begin to characterize the patient mutation, we generated yeast Aac2 L155F. Due to its substantial girth, the insertion of Phe in this pocket was predicted to extrude CL due to steric hindrance (Fig. 5A). This prediction was supported as apo Aac2 was the most dominant population for Aac2 L155F (Fig. 5B), consistent with the corresponding Glu mutant (Fig. 2D, 5B). Since the 2- and 3-CL bound forms were comparatively retained (Fig. 5B), it is likely that the pathogenic Phe residue has a milder impact on pocket 2’s CL association than Glu or Asp residues have. Aac2 L155F was expressed at a similar level to WT (Fig. 5C) and maintained growth capacity in fermentable and respiratory media like L155E (Fig. 5D). The monomeric structure of Aac2 L155F was destabilized on blue-native PAGE and rescued by pre-treatment with CATR or BKA (Fig. 5E). Moreover, Aac2 L155F had significantly reduced ADP/ATP exchange capacity (Fig. 5F-H). These results demonstrate that the patient mutant failed to support Aac2 monomeric structure and transport activity likely due to a perturbed lipid-protein interaction.

**Fig. 5.**
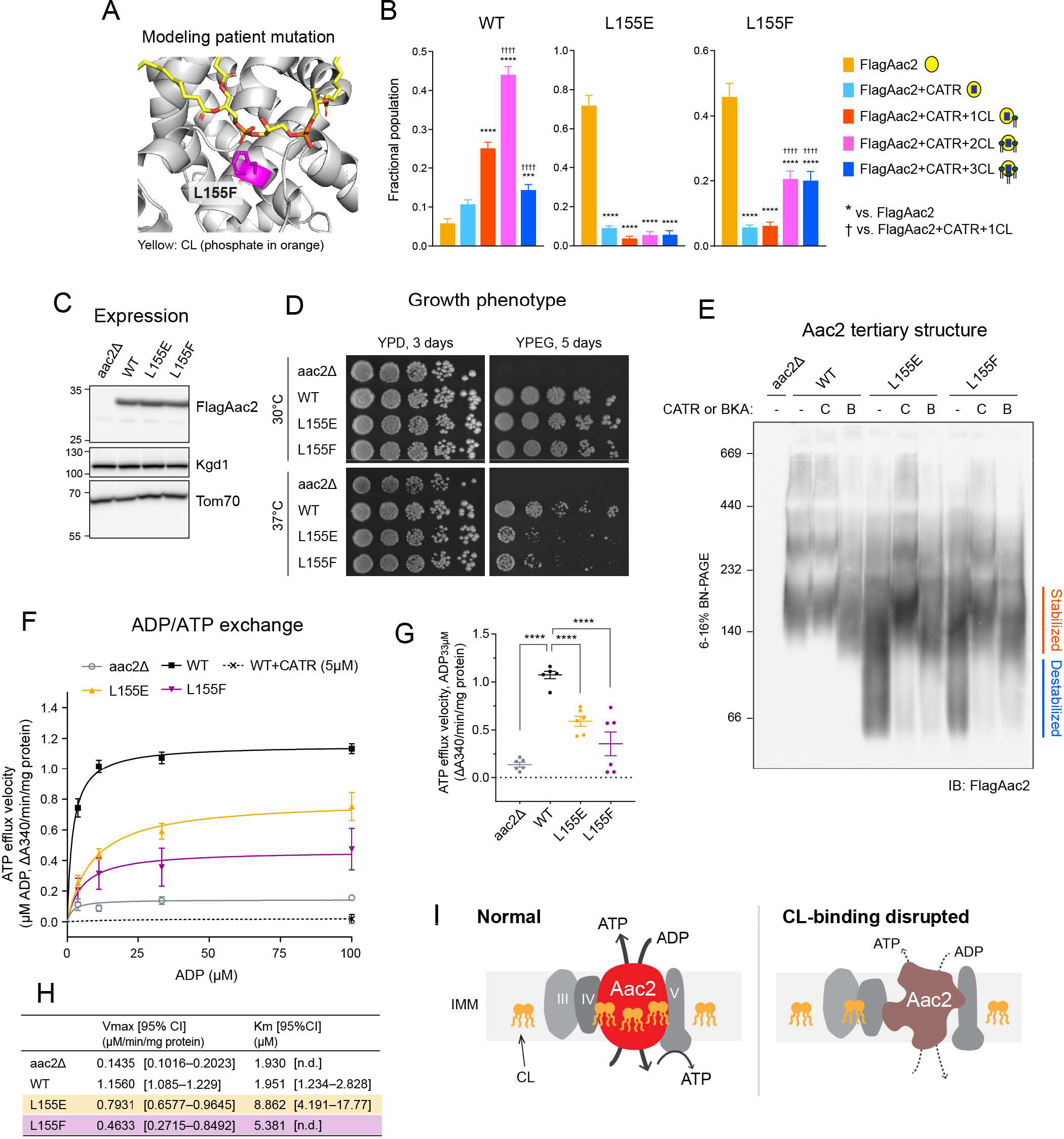
Disturbed Aac2-CL interaction is a pathological mechanism. (A) Mitochondrial myopathy patient mutation L155F was introduced into yeast Aac2 as Fig. 1A. (B) Native MS analysis obtained the fractional population of WT and indicated FlagAac2 mutants associated with CATR and one to three CL molecules. (C) Expression of WT and mutant Aac2 was detected in whole cell extract by immunoblot. Kgd1 and Tom70 served as loading controls (n=4). (D) Growth phenotype of Aac2 CL-binding mutants. Serial dilutions of indicated cells were spotted onto fermentable (YPD) and respiratory (YPEG) media and incubated at 30 or 37°C for 3-5 days (n=4). (E) Aac2 monomeric structure: 100 μg of mitochondria from WT and Aac2-binding mutants were mock-treated or instead incubated with either 40 μM CATR (C) or 10 μM BKA (B) and then solubilized with 1.5% (w/v) digitonin. The extracts were resolved by 6 to 16% blue native-PAGE and immunoblotted for FlagAac2 (n=6). (F-H) ADP/ATP exchange: The efflux of matrix ATP was detected with isolated mitochondria as NADPH formation (A340; absorbance at 340 nm) occurring coupled with *in vitro* glycolysis reaction as Fig. 4. The measurement was performed in the presence of 5 mM malate and 5 mM pyruvate (n=6). (F) The linear part of the initial velocity following the addition of ADP at indicated concentrations was plotted (mean with SEM). Curve fitting was performed by nonlinear regression. (G) The linear part of velocity when 33 μM ADP was added shown as scatter plots (mean with SEM). (H) Fitted Km and Vmax values were obtained by the Michaelis-Menten equation from the replicated experiments (mean). Significant differences were obtained by one-way ANOVA with Dunnett’s multiple comparisons test (vs. WT); *p<0.05, **p<0.01, ***p<0.001, ****p<0.0001. (I) Predicted roles of the buried CLs within Aac2. If CLs are dissociated, Aac2 monomeric structure is destabilized and ADP/ATP transport is compromised, which disrupts energy production via OXPHOS. In C-E, representative images from the indicated replicates are shown.

We sought to extend our investigation of the patient mutation into human ANT1 as expressed in a human cell model. To this end, we took advantage of ant^null^ 293 cells in which all three major ANT isoforms were knocked out (fig. S10) and re-introduced WT and the mutant ANT1 alleles (L141E and L141F). As in yeast Aac2, the introduced L141E and L141F mutations in human ANT1 were predicted to dismiss CL from the peptide backbone (Fig. 6A): L141E could repulse the CL phosphate and L141F sterically block CL engagement. WT and mutant ANT1 were equally expressed (Fig. 6B). As determined by blue-native PAGE, WT ANT1 migrated around 140 kDa, which based on the yeast model, is predicted to reflect a stabilized monomer (Fig. 6C). Pretreatments with CATR and BKA stabilized WT ANT1 at slightly different sizes (Fig. 6C); this migration pattern was identical to yeast Aac2 and likely reflects the conformational status of ANT1 (c-state and m-state) ^35^. Intriguingly, ANT1 L141E and L141F migrated as destabilized smears (Fig. 6C), suggesting that CL-binding to ANT1 in the vicinity of L141 (pocket 2) plays a significant role in stabilizing the carrier’s structure. CATR or BKA pretreatment preserved the structure of L141E and L141F, indicating that the mutant forms of ANT1 were properly folded within the membrane pre-solubilization. The ADP/ATP exchange capacities of L141E and L141F were significantly impaired (Fig. 6D, E). Oxygen consumption measured in cells expressing mutant ANT1 showed compromised basal and ATP production-coupled respiration for L141E and L141F and reduced maximal respiration for L141E-expressing cells (Fig. 6F).

**Fig. 6.**
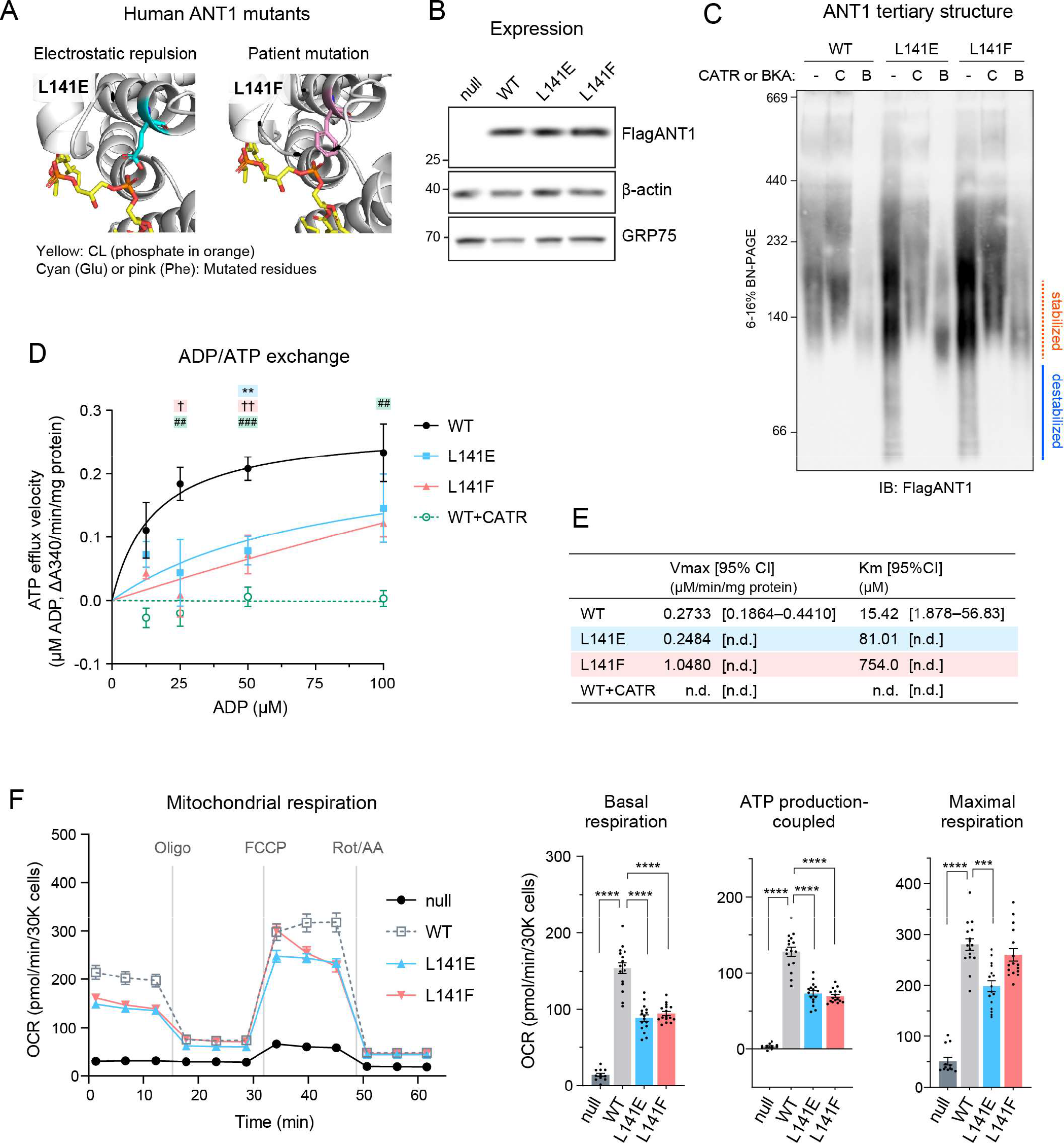
Human ANT1 L141F are structurally and functionally compromised. (A) Human ANT1 was modeled onto bovine ANT1 (PDB ID: 2C3E) using SWISS-MODEL. (B-F) FlagANT1 and the indicated ANT1 mutants were induced by 0.25 μg/ml doxycycline in ant^null^ T-REx-293 cells with three ANT isoforms (ANT1, 2, and 3) knocked out. (B) Expression of ANT1 was detected in whole cell extracts by immunoblot. β-actin and GRP75 were loading controls (n=3). (C) ANT1 monomeric structure: 80 μg of mitochondria from WT and ANT1 mutants were mock-treated or instead incubated with either 40 μM CATR or 10 μM BKA and then solubilized with 1.5% (w/v) digitonin. The extracts were resolved by 6 to 16% blue native-PAGE and immunoblotted for FlagANT1 (n=3). (D-E) ADP/ATP exchange: The efflux of matrix ATP was detected with isolated mitochondria as NADPH formation (A340; absorbance at 340 nm) as Fig. 4. The measurement was performed in the presence of 5 mM malate and 5 mM pyruvate. 5 μM CATR was added to WT mitochondria prior to stimulating the efflux (n=3). (D) The initial velocity following the addition of ADP at indicated concentrations was plotted (mean with SEM). Curve fitting was performed by nonlinear regression. Significant differences obtained by one-way ANOVA with Dunnett’s multiple comparisons test are shown as *, L141E; †, L141F; #, WT+CATR (vs. WT). *p<0.05, **p<0.01, ***p<0.001. (E) Fitted Km and Vmax values from the Michaelis-Menten equation (mean). (F) Cellular oxygen consumption rate (OCR) was measured using a Seahorse XF96e Flux Analyzer with the Mito Stress Test kit under indicated conditions. Basal and maximal OCR were obtained under glucose stimulation after FCCP treatment to uncouple mitochondria. ATP production-coupled respiration is defined as basal OCR subtracted by post-oligomycin OCR. Significant differences were determined by one-way ANOVA with Dunnett’s multiple comparisons test (vs. WT), ***p<0.001 ****p < 0.0001. Means with SEM (n=16). In B and C, representative images from the indicated replicates are shown.

To investigate the impact of L141 mutations on the molecular structure and CL interactions of ANT1, all-atom MD simulations of WT and mutant human ANT1 were performed in a CL containing lipid bilayer (fig. S11). Time-averaged probability density maps of CL positions relative to ANT1 were generated for the “CL prebound” (Fig. 7A) and “CL unbound” (Fig. 7B) simulations. The “CL prebound” and “CL unbound” simulations are defined by the presence (prebound) or absence (unbound) of a CL lipid in the vicinity of binding pocket 2 prior to the initial state of the simulations (See Methods and figs. S11B-C for further details). For the prebound simulations for WT and mutant ANT1, we observed a high density of CL (Fig. 7A) in the three known CL-specific binding pockets^45–48^. In the prebound simulations, we do not observe a significant alteration to the CL densities around pocket 2. This may suggest longer time scales are required to capture the complete binding and unbinding processes of CL from the binding pockets. Another factor, in the case of the L141E mutant, is that the expected electrostatic repulsion between CL and the Glu residue is mitigated by formation of a salt bridge interaction with the proximal R152 residue.

**Fig. 7:**
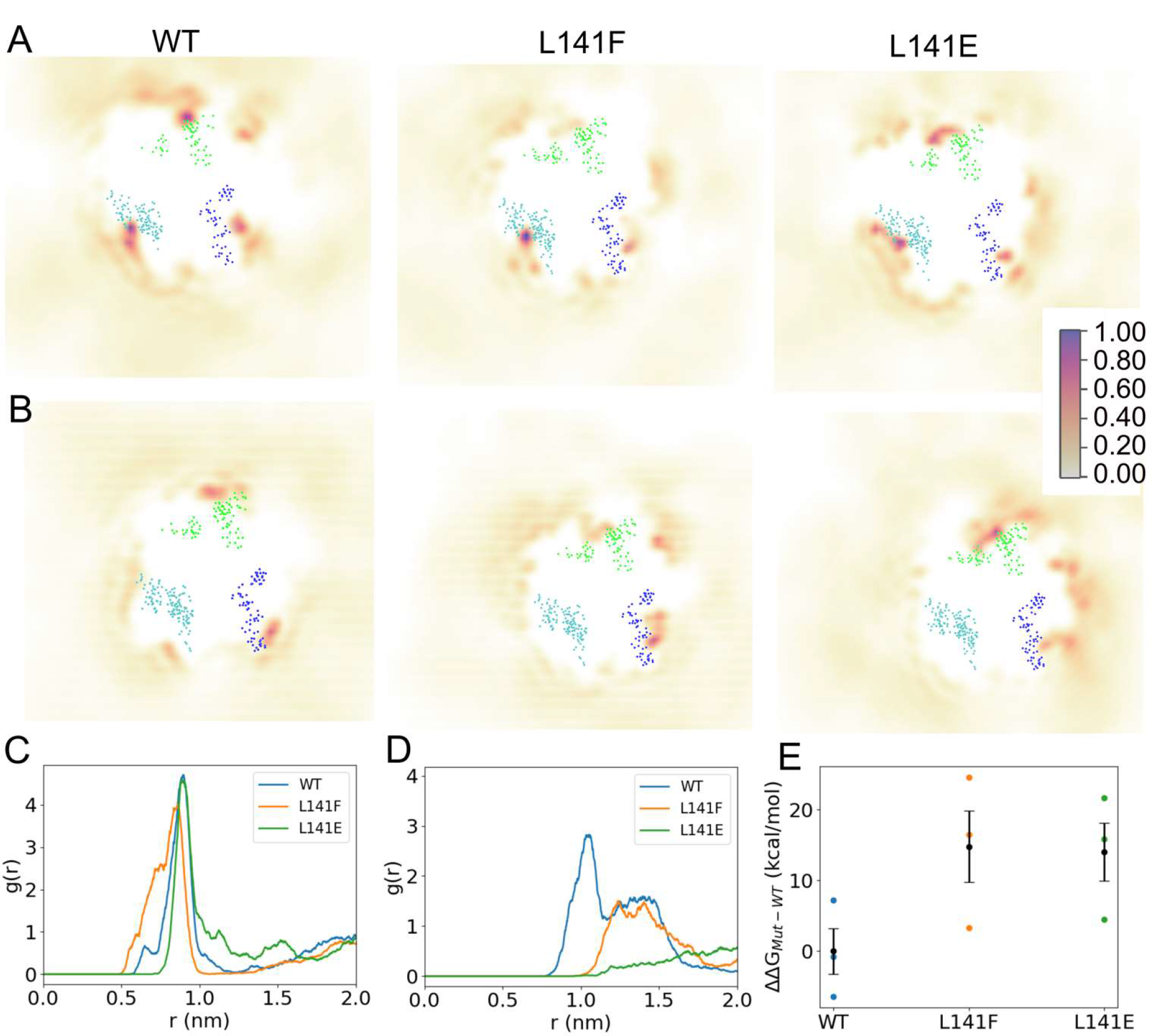
**MD simulations predict reduced CL-ANT1 affinity at pocket 2 for L141 mutants**. (A-B). Time-averaged 2D density maps of CL lipids for the equilibrium prebound (A) and unbound (B) simulations. The CL binding pockets are defined by the blue (pocket 1), cyan (pocket 2) and green (pocket 3) dots, which represent the amino acids that comprise each binding site. The density surfaces were calculated using vmd volmap with 1 Å resolution. Occupancy scale bar on right applies to all images. (C-D) Radial distribution function profile for the identification of CL lipids around residue 141 for prebound (C) and unbound (D) simulation. (E) The estimated relative free energy required to decouple CL from binding pocket 2 of ANT1 for WT and mutants L141F and L141E. Using Wilcoxson rank sum test between WT and L141E indicates statistical significance (p< 0.05), while the difference between WT and L141F is not significant. **Note:** Binding site residues considered in the present study are based on the protein-CL lipid interactions and are the selected amino acid residues within 8 Å of the bound CL molecules: **Pocket 1:** 36, 53, 54, 55, 271, 272, 273, 274, 275, and 276; **Pocket 2:** 71, 72, 73, 74, 75, 141, 152, 155, 156, 157, and 158**; Pocket 3:** 251, 252, 253, 254, 255, 174, 175, 176, 177, and 178, respectively.

As an alternative to extending the simulation time, we performed a set of simulations in which the CL lipids were removed from pocket 2 (unbound). From these simulations, we observed for WT ANT1 a density map consistent with the prebound simulations showing a high density of CL located in the three known binding sites^45–48^. In contrast, the mutant ANT1 systems displayed low CL density proximal to pocket 2 (Fig. 7B). The 2D lipid density profiles are supported by 2D radial distribution functions (g(r)), which represent the probability of locating the central glycerol of CL at varying radial distances from the Cα atom of residue 141. In the case of prebound simulations (Fig. 7C), both WT and mutant proteins had the highest probability of finding CL lipids to be at ∼0.90 nm from residue 141, which is consistent with crystallographically determined distances (PDBID 1OKC: *r141-CL*=0.94 nm; 2C3E: *r141-CL*=0.96 nm). In the case of the unbound simulations (Fig. 7D), a considerably higher probability of CL lipids proximal to residue 141 was observed for WT compared to L141F or L141E mutant systems. Both WT and L141F displayed a plateau region centered around 1.3 nm, which may represent an approach state prior to CL accommodation in pocket 2. The bulky Phe residue of L141F may impede the accommodation of CL into pocket 2 and the lack of the approach plateau in L141E is likely due to the electrostatic repulsion between Glu and CL. Unlike the prebound simulations, a salt bridge between E141 and R152 is not observed for the L141E system in the unbound simulations. It is possible that the presence of CL in pocket 2 in the prebound simulations drove the formation of the E141-R152 salt bridge, and when CL is removed from the pocket (unbound) the formation of the salt bridge is less energetically favorable.

Protein structure and dynamics during the simulations were examined and ANT1 was more stable in the CL prebound simulations than the CL unbound simulations, for WT and mutants (figs. S12A-B). To examine the dynamics of residue 141 and the stability of pocket 2 residues, the distance between the Cα atom of residue 141 with that of the other pocket 2 binding residues was calculated for the prebound simulations (fig. S12C). In the WT system, the arrangement of pocket 2 residues relative to L141 was highly stable. On the contrary, neighboring amino acids of residue 141 such as residues 152 and 155-158, displayed fluctuating distances, especially His155 in the L141F mutant system. The observed shorter distances between H155 and F141 in L141F indicate instances of π − π stacking interactions. The L141E mutant system showed comparatively less variable distances between residue 141 and all other considered pocket 2 binding residues than the L141F mutant. This is potentially due to the observed salt bridge interaction between residue E141 and R152. Overall, the simulations support that mutations of L141 destabilize the CL binding environment and cause higher fluctuations in the residues presented between helices 3-4 (matrix-oriented loop).

Assimilating the prebound (Fig. 7A, C) and unbound (Fig. 7B, D) simulation results, the CL interaction at pocket 2 is stable in both WT and mutants, but the mutations may create a barrier for CL to access the binding site, resulting in the reduced pocket 2 density in the mutants. Note that the equilibrium simulations are not sufficiently long to capture unbinding/rebinding events to accurately calculate populations (i.e. free energy differences) and therefore we used a free energy perturbation (FEP) approach to estimate the difference in CL binding free energies between WT and mutants (fig. S13). The relative free energy required to decouple a CL molecule from pocket 2 of ANT1 WT, L141F, and L141E systems are shown in Fig. 7E. From the calculated *ΔΔG* values, WT ANT1 showed ∼12 kcal/mol more favorable binding energy than the two studied mutant systems. *In toto*, these combined experimental and simulation results demonstrate that CL-binding at pocket 2 of human ANT1 is both structurally and functionally important and that the mitochondrial dysfunction associated with the previously uncharacterized L141F pathogenic mutation ^44^ derives from a perturbed CL-ANT1 interaction.

## Discussion

Previous structural studies identified three CL molecules tightly bound to AAC in an evolutionally conserved manner ^7, 8^; however, the functional significance of these lipid-protein interactions has not been determined. In this study, we engineered yeast Aac2 mutants to disrupt CL binding; the dissociated lipid-protein interactions of the mutants were confirmed by native MS analysis (Fig. 2). The results provide experimental evidence that the CL molecules bound to Aac2 stabilize the carrier’s monomeric structure and support its transport activity (Fig. 5I). Further, we identified that the residues mutated to disrupt CL-Aac2 interactions are conserved in mammalian ANTs, which includes a previously reported patient mutation (L141F in human ANT1) ^44^. When the patient mutation was modeled in yeast Aac2 (L155F) and human ANT1 (L141F), the pathogenic mutation compromised the carrier’s structure and transport activity (Fig. 5, Fig. 6) due to a disturbed lipid-protein interaction (Fig. 5B, Fig 7). This would represent, to the best of our knowledge, the first known disease-causing mutation which disrupts a structurally and functionally important lipid-protein interaction.

The patient mutant, yeast Aac2 L155F, somewhat retained its capacity to bind 2-3 CLs when assessed by our native MS system, even though apo Aac2 was the most dominant population for this mutant similar to the negatively charged residue mutants (Fig. 2D, 5B). This suggests that the Phe residue has a relatively mild effect on dissociating the targeted CL-Aac2 interaction compared to the electrostatic repulsion between CL phosphates and introduced negatively charged amino acids in the Glu or Asp mutants. Our MD simulations further clarified the CL-binding status of the patient mutant ANT1 L141F (Fig. 7). The unbound simulation indicated that L141F failed to accommodate CL in pocket 2 but retained a CL approaching event, in contrast to L141E which lost both events (Fig. 7D). Nonetheless, the extent of structural and functional impairments was comparable between the Glu and Phe mutants in yeast Aac2 and human ANT1 (Fig. 5, Fig. 6). These results imply that the disruption of this CL-Aac2 interaction, even if incomplete, causes serious impairment.

Our results indicate that the CL molecules in each pocket behave distinctly. Whereas the mutants for pocket 2 (N90E, L155E, G172E) and pocket 3 (R191D, M255E, G267E) had severely compromised transport as well as OXPHOS capacity, the pocket 1 mutants (I51E and G69D) largely retained these activities. The double mutant (G69D/G267E) for pockets 1 and 3 showed a stronger defect in transport capacity than the corresponding single mutants. Our native MS results indicated that the strength of CL bindings is different among the pockets: one of the pockets has a relatively weak affinity and the other two pockets have a similar affinity (fig. S2). In repeated experiments to obtain fractional populations using distinct preparations of mitochondria, we noted that the preferred interaction status for WT Aac2 varied slightly: CATR- bound Aac2 + 3 CLs was the highest in Fig. 2B whereas CATR-bound Aac2 + 2 CLs was the highest in Fig. 5B. While the basis for this difference is unclear, we note that even when Aac2 + 2 CLs was dominant, a significant amount of Aac2 + 3 CLs remained. In the net, these data indicate that WT Aac2 binds up to three CLs and also strongly suggest that one of the three CL interactions engages Aac2 with a lower affinity than the other two CLs. Although we expected that fractional population results of the mutants from the native MS analyses might give us insight into the CL-binding affinity of each pocket, the broad impact on CL binding of most mutants made this difficult. Recent MD simulations of fungal AAC in the m-state demonstrated that CLs in pockets 2 and 3 (CDL800 and CDL802) are constantly bound while CL in pocket 1 (CDL801) is more labile ^33^. Supporting our MSMS results, these simulation results suggest that the functional difference across pockets that we observed may correlate with their binding affinity for CLs. Another MD simulation of bovine ANT1 in the c-state indicated that pocket 2 is densely associated with CL and this CL contributes to stabilizing the carrier ^34^. Collectively, pockets 2 and 3 appear to have preeminent roles while the labile CL molecule in pocket 1 would have a more nuanced and/or specific Aac2-related role. Although we performed MD simulations only on the pocket 2 mutant of human ANT1, the simulations will be extended to the other CL-binding pocket mutants with physiological CL amount and acyl chain profile (e.g. representing the human heart), which would further clarify the binding affinity across pockets and provide insight into the physiological significance of these specific lipid interaction events.

Worth considering is that some of the inserted mutations are likely to exert non-CL binding related effects. For instance, N90E and R191D are located in the vicinity of several residues recently shown to be critical for substrate binding related to the AAC transport cycle ^49^. It would seem plausible that the negatively charged residues we inserted might affect substrate binding itself. Indeed, these two mutants had severe respiratory growth defects (Fig. 1C). In addition, the N90E mutant had a weakened association with respiratory complex subunits compared to the other mutants tested (fig. S8). Still, CL binding was disrupted for both the N90E and R191D Aac2 mutants (Fig. 2D). As such, their defects may be attributed to a combination of disturbed lipid-protein interaction and impaired substrate binding, and both may be affected in a distinct and/or synergized manner.

Most of the carriers belonging to the SLC25 family are embedded in the IMM like AAC. Their topology and transport-related conformations have been predicted to be very similar to AAC ^50^. Given that CL is enriched in the IMM, CL may be widely involved in supporting the structure and transport activity of the extended SLC25 family. In fact, a very recent study indicated that CL binding is involved in the activity of SLC25A51, also known as the mitochondrial NAD^+^ carrier^51^. Our findings highlight the structural and functional significance of tightly-bound CL in yeast Aac2 and human ANT1. This serves as a paradigm for the important roles exerted by specific lipid-protein interactions in mitochondria and provides a clinically relevant example of conserved lipid-protein interactions in a membrane-integrated carrier protein.

## Limitations of this study

Although our rationale for inserting Glu or Asp into the Aac2 mutants is based on the logical expectation that electrostatic repulsion between acidic residues and CL phosphates would prevent tight CL binding, it is also possible that these residues may alter the local secondary structure which in turn blocks CL binding. Another limitation is that the measurements of AAC/ANT transport activity performed in this study rely on the function of the electron transport chain. Based on CCCP-induced respiration rates (i.e. uncoupled respiration from ADP phosphorylation) (fig. S5), expression of mitochondrial DNA-encoded CIV subunits (fig. S6), and Aac2-RSC interactions (fig. S7, 8), Aac2 pocket 2 mutants have measurable electron transport chain defects that could contribute to the severely decreased transport activity observed for this subset of mutants. As the possibility that these defects could impact AAC transport activity cannot be excluded, further investigation may be required to clarify the activity intrinsic to AAC/ANT using *in vitro* systems and pharmacologically generated membrane potentials.

## Materials and Methods

### Yeast strains and growth conditions

All yeast strains used were derived from *S. cerevisiae* parental strain GA74-1A (*MATa, his3- 11,15, leu2, ura3, trp1, ade8, rho+, and mit+*). *aac2*Δ were generated by replacing the entire open reading frame of AAC2 with markers HIS3MX6, as previously described ^24^ via polymerase chain reaction (PCR)–mediated gene replacement ^52^.

Single point mutations into Aac2 open reading frame were introduced by PCR-based overlap extension using primers containing each mutant sequence. To place Flag tag onto the N-terminus of Aac2 still downstream of the promoter, PCR-mediated overlap extension was performed against either WT Aac2 or each mutant open reading frame using primers containing Flag sequence (DYKDDDDK). All were cloned into pRS305. Primers used to generate the constructs are listed in Table S1. The sequences of every construct were verified by Sanger DNA sequencing. The pRS305 constructs were linearized and integrated into the *LEU* locus in the *aac2*Δ background. Clones were selected on synthetic dropout medium (0.17% (w/v) yeast nitrogen base, 0.5% (w/v) ammonium sulfate, 0.2% (w/v) dropout mixture synthetic-leu, 2% (w/v) dextrose) and verified by immunoblot.

*crd1*Δ strain was generated by homology-integrated CRISPR-Cas system as previously described ^53–55^. CRISPR-Cas9 gene block was designed to target *CRD1* and cloned into pCRCT plasmid ^53^. pCRCT was a gift from Huimin Zhao (Addgene plasmid #60621; http://n2t.net/addgene:60621; RRID: Addgene_60621). The sequence of *CRD1* gene block is: 5’- CTTTGGTCTCACCAAAACGATGTCCTAGAAAGAGGAGATGAATTTTTAGAAGCCTATCCCAGAAGAAGGCAGATTTCTTTCATACAGTACATCATTACTGACCTTCGGTGTATCAAAAGTATGAAAGAAATCTCCAAAGTTTTAGAGAGAGACCTTTC-3’.

To map the epitope(s) recognized by the in-house generated ANT2 antiserum ^56^, we subcloned WT ANT3, ANT2, or the indicated ANT2 epitope mutants in which ANT2-specific residues were replaced by their ANT3 counterparts by overlap extension PCR, into pRS315 (fig. S10A, B). Of note, functional expression of human ANT isoforms in yeast requires the substitution of a stretch of mammalian N-terminal amino acids for those from yeast Aac2 ^41^, which allows their detection with the Aac2-specific monoclonal antibody, 6H8 ^36^. All constructs were heterologously expressed in *aac2*Δ yeast and grown in synthetic dropout media (0.17% yeast nitrogen base, 0.5% ammonium sulfate, 0.2% dropout mix synthetic –leu) supplemented with 2% dextrose.

Yeast were grown in nutrient-rich YP [1% (w/v) yeast extract and 2% (w/v) tryptone] media that contained 2% (w/v) dextrose (YPD). For growth analysis on solid plates, yeast cells were grown in YPD before spotting onto YPD or YPEG [1% (w/v) yeast extract, 2% (w/v) tryptone, 1% (v/v) ethanol, and 3% (v/v) glycerol] plates [2% (w/v) Bacto agar]. For mitochondrial isolation, cells were grown in YP-Sucrose [1% (w/v) yeast extract and 2% (w/v) tryptone, 2% (w/v) sucrose].

### Human cell lines and culture conditions

T-REx™ 293 cells (Invitrogen R71007) were grown on a plate coated by Cultrex Rat Collagen I (50 µg/ml, R&D systems 3440-100-01) at 37°C in 5% CO2 in high glucose-containing DMEM (Corning 10-013-CM) supplemented with 10% FBS (CPS Serum FBS-500), 2 mM L-glutamine (Gibco 25030081), 1 mM sodium pyruvate (Sigma S8636), 50 µg/ml uridine (Sigma U3003), and 15 µg/ml blasticidin (InvivoGen ant-bl-1). We have ruled out the presence of mycoplasma in the cells through routine testing.

ant^null^ cells were established by knocking out ANT1, ANT2, and ANT3 in T-REx™ 293 cells. Individual gRNAs that targeted ANT1, ANT2, and ANT3 were separately cloned into pSpCas9(BB)-2A-Puro (PX459) V2.0, a gift from Feng Zhang (Addgene plasmid #62988; http://n2t.net/addgene:62988; RRID: Addgene_62988)^57^. gRNA target sequences are 5’- GATGGGCGCTACCGCCGTCT-3’ for ANT1, 5’-GGGGAAGTAACGGATCACGT-3’ for ANT2, and 5’-CGGCCGTGGCTCCGATCGAG-3’ for ANT3, respectively. Transfections of the PX459 constructs were performed individually using FuGENE 6 Transfection Reagent (Promega E2691) according to the manufacturer’s protocol. Following each transfection, cells were selected with puromycin (2 µg/ml) for 72 hours, single clones were isolated by ring cloning, the absence of the protein expression was analyzed by immunoblotting, and the next transfection proceeded.

Flag-ANT1 was amplified by PCR using the previously published construct ^58^ as a template with primers designed to attach the Flag-tag onto the N-terminus of ANT1. Single point mutation into ANT1 open reading frame was introduced by PCR-based overlap extension using primers containing each mutant sequence. Table S2 lists sequences of the primers used for these constructs. All were cloned into pcDNA™5/FRT/TO (Invitrogen V652020). ant^null^ cells were co-transfected with pOG44 (Invitrogen V600520, expressing the Flp-recombinase) and the relevant pcDNA™5/FRT/TO plasmid at a ratio of 9:1 using FuGENE 6 Transfection Reagent. Transfected cells were selected using hygromycin B (Invitrogen) at 20 µg/ml for 2 days and then 40 µg/ml for 4 days. Individual clones isolated by single-cell seeding on a 96-well plate were subsequently expanded and screened by immunoblot. The cells introduced with Flag-ANT1 WT and mutants were maintained in glucose-based media [high glucose DMEM, 10% FBS, 2 mM L-glutamine, 1 mM sodium pyruvate, 50 µg/ml uridine, 15 µg/ml blasticidin] supplemented with 0.25 µg/ml doxycycline.

### Whole-cell protein extraction and immunoblotting

Protein extraction from yeast cells and immunoblotting were performed as detailed previously ^24, 55, 59^. For human cells, proteins were extracted from confluent six- or twelve-well tissue culture dishes using RIPA lysis buffer [1% (v/v) Triton X-100, 20 mM HEPES-KOH, pH 7.4, 50 mM NaCl, 1 mM EDTA, 2.5 mM MgCl2, 0.1% (w/v) SDS] spiked with 1 mM PMSF and quantified using the bicinchoninic acid assay (Pierce) as described in ^60^.

Antibodies used in this study are listed in Table S3. Custom-produced antibodies against yeast Tim54 were generated by Pacific Immunology (Ramona, CA) using affinity purified His6Tim54 as antigen. In brief, the predicted mature Tim54 open reading frame starting at Lys-15 was cloned downstream of the His6 tag encoded in the pET28a plasmid (Novogene) and transformed in BL21(RIL) *Escherichia coli*. A 1L culture was induced with 0.5 mM IPTG at 30°C for 4 hrs and the collected bacterial pellet (3020 x *g* for 10 min) was washed with 0.9% (w/v) NaCl and stored at -20°C until purification was performed. The bacterial pellet was resuspended in 40 ml lysis buffer (50 mM NaH2PO4, 300 mM NaCl, 10 mM Imidazole, 0.1 mM EDTA, pH 8.0), lysozyme added for 1 mg/ml, and incubated with rocking for 30 min at 4°C. The bacterial suspension was ruptured using an Avestin Homogenizer and the resulting lysate centrifuged at 10,000 x *g* at 4°C for 20 min. The pellet was solubilized with freshly made Inclusion Body solubilization buffer (1.67%(w/v) Sarkosyl, 10 mM DTT, 10 mM Tris-Cl pH 7.4, 0.05% (w/v) PEG3350) by vortexing on high and then incubated on ice for 20 min. 10 ml of 10 mM Tris-Cl pH 7.4 was added to the suspension which was then centrifuged at 12,000 x *g* at 4°C for 10 min. The recovered material was subjected to ammonium sulfate precipitation, with the recombinant Tim54 crashing out of solution at 20% (w/v) ammonium sulfate, and the resulting pellet was resuspended in 2 ml of Inclusion Body solubilization buffer base (10 mM DTT, 10 mM Tris-Cl pH 7.4, 0.05% (w/v) PEG3350) containing 0.5%(w/v) Sarkosyl. To this suspension, 8 ml of Wash Buffer A (0.1% (w/v) Sarkosyl, 50 mM NaH2PO4, 300 mM NaCl, 20mM Imidazole, 10% (v/v) glycerol, pH 8.0) was added and the solution nutated for 30 min at room temperature. His6Tim54 was then purified from this suspension by incubating with 3 ml Ni-NTA in a capped column rotating for 2 hrs at room temperature. Following two column-volume washes with 1) 0.1% (w/v) Sarkosyl, 50 mM NaH2PO4, 300 mM NaCl, 20 mM Imidazole, 10% (v/v) glycerol, pH 8.0; and 2) 0.1% (w/v) Sarkosyl, 50 mM NaH2PO4, 600 mM NaCl, 20 mM Imidazole, 10% (v/v) glycerol, pH 8.0, bound material was recovered with elution buffer (250 mM imidazole, 0.1% (w/v) Sarkosyl, 50 mM NaH2PO4, 300 mM NaCl, and 10% glycerol, pH 8.0; 6 sequential 0.5 ml elutions). Protein-containing fractions, identified using the Bradford Assay (Bio-Rad), were combined, PBS dialyzed, and quantified using a BSA standard curve prior to antibody generation.

### Mitochondrial isolation

Isolation of mitochondria from yeast cells was performed as previously described ^55^. For mitochondrial isolation from T-REx-293 cells, cells were seeded onto three or more 150 mm × 25 mm tissue-culture dishes and allowed to expand in glucose-based medium supplemented with 0.25 μg/ml doxycycline. 48 hrs before isolation, the cells that reached 70-80% confluency were fed fresh medium containing 0.25 μg/ml doxycycline. Mitochondrial isolation was performed according to a previous protocol ^61^. Briefly, cells were homogenized with IBc buffer [200 mM sucrose, 10 mM Tris-MOPS, 1 mM EGTA/Tris, pH 7.4)] using a Teflon Potter-Elvehjem motor-driven dounce set at 1,600 rpm. Homogenates were centrifuged twice at 600 g for 10 min at 4°C to precipitate debris and the nuclear fraction. The supernatant was centrifuged at 7,000 g for 10 min at 4°C. The resulting pellet was resuspended in IBc buffer and centrifuged at 7,000 g for 10 min at 4°C. This was repeated at 10,000 g for 10 min at 4°C. The final pellet was resuspended in IBc buffer. If not used immediately, pellets were aliquoted, snap frozen with liquid N2, and stored at −80°C for downstream analyses.

### Aac2 membrane topology

50 μg of intact mitochondria (in 1 ml BB7.4 [0.6 M sorbitol, 20 mM K+ HEPES (pH7.4)]) or mitoplast were incubated with or without proteinase K (40 μg/ml) on ice. Mitoplasts were obtained by osmotic swelling; to mitochondria suspended in 0.05 ml BB7.4, 19X volumes of 20 mM K+ HEPES, pH7.4 (total 1 ml) was added. After 30 min incubation on ice, 5 mM PMSF was added to deactivate proteinase K. The samples were precipitated by TCA, heated at 60 °C for 5 min, and kept on ice for 5 min for recovery. Following centrifugation at 21,000 g for 20 min at 4 °C, the pellet was washed with cold acetone and resuspended in 0.1M NaOH. An equal volume of 2X reducing buffer was added and then boiled for 5 min at 95°C. The samples were analyzed by SDS-PAGE followed by immunoblotting.

### Purification of Aac2 for native MS analysis

8 mg mitochondrial membranes were thawed on ice and reconstituted in BB7.4 (0.6M sorbitol in 20 mM HEPES buffer) containing 40 μM CATR (Sigma C4992) for 20 min at 4°C. After centrifuging at 21,000 g, the membrane pellets were solubilized in 1 ml lysis buffer (20 mM Tris pH 8.0, 150 mM NaCl, 10% glycerol, protease inhibitor cocktail, 2% UDM (Anatrace U300LA) on a rotary shaker for 30 min. Solubilized mitochondrial proteins were obtained by centrifuging at 21,000g for 30 min at 4°C. Anti-Flag magnetic beads were equilibrated with lysis buffer by washing twice and incubated with the solubilized membrane protein for 2 hrs on a rotary shaker at 4°C. The lysate was then aspirated and washed once with lysis buffer and twice with wash buffer (20 mM Tris, pH 8.0, 150 mM NaCl, 10% glycerol, protease inhibitor cocktail, 0.06% UDM). Flag-tagged Aac2 was eluted by incubating with 1 ml elution buffer (20 mM Tris pH 8.0, 150 mM NaCl, 10% glycerol, 0.6% UDM, 125 μg Flag peptide) for 1 hr in a rotary shaker at 4°C. Flag- Aac2 was concentrated to 5 μM and stored in -80°C until further use for MS analysis. Purifications were done from two different preparations, with three biological replicates.

### Native MS

Prior to MS analysis, UDM solubilized Flag-Aac2 were buffer-exchanged into 200 mM ammonium acetate at pH 8.0, with 2 × CMC (critical micelle concentration) of the LDAO (Anatrace D360) detergent using a 7 kDa Zeba spin desalting columns (Thermo). The LDAO exchanged Flag-Aac2 was then introduced directly into the mass spectrometer using gold-coated capillary needles (prepared in-house). Data were collected on a UHMR mass spectrometer (Thermo Fisher Scientific) optimized for analysis of high-mass complexes, using methods previously described ^62^. The instrument parameters used were as follows: Positive polarity, capillary voltage 1.2 kV, quadrupole selection from 1,000 to 15,000 m/z range, S-lens RF 100%, argon UHV pressure 1.12 × 10^-9^ mbar, temperature 200 °C, resolution of the instrument 17,500 at m/z = 200 (a transient time of 64 ms) and ion transfer optics (injection flatapole, inter-flatapole lens, bent flatapole, transfer multipole: 8, 7, 6 and 4 V, respectively). The noise level was set at 3 rather than the default value of 4.64. In-source trapping of -150 eV was used to release the protein out of the detergent micelles. No collisional activation in the HCD cell was applied at any stage. All other data were visualized and exported using Xcalibur 4.1.31.9 (Thermo Scientific). The relative intensities of apo, CATR, and CL bound species were obtained by deconvoluting the native MS data using UniDec ^63^. The intensities were converted to mole fractions to determine the fractional populations of each ion in the spectra. Similar parameters were used for data processing in UniDec when comparisons are made. For MSMS experiments, the highest intensity charge state was selected in the quadrupole and subjected to HCD from 50-300V. Each biological replicate was analyzed with at least two technical replicates.

PE (palmitoyl-oleoyl; Anatrace P416), PG (palmitoyl-oleoyl; Anatrace P616), lyso-PC (palmitoyl; Avanti 855675) and CL (tetramyristoyl; Avanti 710332) were used for lipid add-back experiments. Phospholipids were prepared according to the method described before ^64^. In brief, stock solutions of 3 mM phospholipids were prepared in 200 mM ammonium acetate and stored in -20°C until use. These were further diluted into 200 mM ammonium acetate containing 2 × CMC of LDAO before titrations and used for further experiments. For titrations, 2.5 μM Flag- Aac2 protein eluted from *crd1*Δ strain were incubated with required amount of lipid at different concentration points and immediately sprayed into the mass spectrometer.

### Blue native-PAGE

One-dimensional (1D) blue native-PAGE was performed as previously described ^59^. Briefly, mitochondria were solubilized in lysis buffer [20 mM Tris-Cl, 10% (v/v) glycerol, 100 mM NaCl, 20 mM imidazole, 1 mM CaCl2 (pH 7.4)] supplemented with protease inhibitors (1 mM PMSF, 2 μM pepstatin A, 10 μM leupeptin) containing either 1.5% (w/v) digitonin (Biosynth D-3200) or 2% (w/v) UDM. For pre-treatment of yeast mitochondria with CATR or BKA, mitochondria were incubated in BB7.4 with 40 μM CATR (Sigma C4992) or BB6.0 [0.6 M sorbitol, 20 mM KOH-MES (pH6.0)] with 10 μM BKA (Sigma B6179) for 15 min on ice as reported previously ^35^, pelleted at 21,000 g for 5 min and then solubilized as above. For human cell mitochondria, IBc pH7.4 for CATR or pH6.0 for BKA was used during incubation. Protein extracts were collected as supernatant following centrifugation (21,000 g for 30 min, 4°C) and mixed with 10X blue native-PAGE sample buffer [5% (w/v) coomassie brilliant blue G-250 (Serva), 0.5 M 6- aminocaproic acid, and 10 mM bis-tris/HCl (pH 7.0)]. The extracts were resolved by 6 to 16% or 5 to 12% house-made 1D blue native-PAGE gels.

### ADP/ATP exchange

ATP efflux from yeast mitochondria was measured according to previous reports ^39–41^ with a slight modification. Isolated mitochondria were suspended in reaction buffer [0.6 M mannitol, 0.1 mM EGTA, 2 mM MgCl2, 10 mM KPi, 10 mM Tris-HCl (pH7.4)] with ATP detection system containing 5 mM α-ketoglutarate (Sigma-Aldrich 75892), 0.01 mM Ap5A (Sigma D6392), 2 mM glucose, hexokinase (2U/reaction, Sigma H4502), glucose-6-phosphate dehydrogenase (2U/reaction, Sigma-Aldrich G5885), and 0.2 mM NADP (Roche 10128031001). When indicated, 5 mM malate (Sigma M1000) and 5 mM pyruvate (Sigma S8636) were added to the reaction buffer. The exchange reaction was performed on a 96-well plate. Following the addition of external ADP (Sigma A2754), NADPH formation was monitored as increasing absorbance at 340 nm using a plate reader. 10 or 20 μg of mitochondria per well were loaded in the presence or absence of malate and pyruvate, respectively. The initial linear part of the reaction curve was used to calculate the ATP efflux velocity. The kinetics were analyzed by the Michaelis-Menten equation using Graphpad Prism.

### Oxygen consumption measurement in yeast mitochondria

OCR was measured with isolated yeast mitochondria using a Seahorse Flux Analyzer. Mitochondria were reconstituted in assay buffer [0.25 M sucrose, 5 mM KH2PO4, 2 mM HEPES, 2.5 mM MgCl2, 2 mM EGTA (pH 7.2)] containing fatty acid-free BSA (0.2%). 50 μl of the reconstituted mitochondria were loaded onto an XF96e assay plate (1.8 μg per well for NADH assay, 3 μg per well for succinate assay) and the assay plate was centrifuged at 2,000 g for 10 min at 15°C to enhance adherence. Then, additional 120 μl of assay buffer per well was carefully layered. The compounds were reconstituted in assay buffer without BSA and loaded into the cartridge ports with the following positions at the indicated final concentrations: port A, 5 mM NADH (Millipore 481913) or 10 mM succinate (Alfa Aesar 41983); port B, 1 mM ADP; port C, 10 μM oligomycin (APExBio C3007); port D, 10 μM CCCP (Sigma-Aldrich C2759). OCR was monitored before the first injection and upon each injection with 3 cycles of 1 min-mixing followed by 3 min-measurement at 37 °C.

### Oxygen consumption measurement in human cells

Cells were seeded onto an XF96 cell culture microplate coated with 0.001% (w/v) poly-L-lysin (Sigma-Aldrich P4707) at 30,000 cells/well and incubated for 48 hrs in glucose-based media supplemented with 0.25 μg/ml doxycycline. On the day of measurement, cells were washed twice with Seahorse XF DMEM Basal Medium supplemented with 10 mM glucose, 2 mM L-glutamine, and 1 mM sodium pyruvate and preincubated for 1 hr in a 37°C humidified CO2-free incubator. OCR was measured by a Seahorse XF96e FluxAnalyzer with the Mito Stress Test kit (Agilent). The Mito Stress Test inhibitors were injected during the measurements as follows; oligomycin (2 μM), FCCP (0.5 μM), rotenone and antimycin A (0.5 μM). The OCR values were normalized to cell density determined by the CyQUANT Cell Proliferation Assay Kit (Invitrogen C7026) according to the manufacturer’s instruction.

### Immunoprecipitation

As indicated above, mitochondria (250 μg) were pre-treated with 40 μM CATR in BB7.4. As performed previously ^35^, the treated mitochondria were pelleted at 21,000 g for 5 min at 4 °C and solubilized with lysis buffer [20 mM Tris-Cl (pH 7.4), 10% (v/v) glycerol, 100 mM NaCl, 20 mM imidazole, 1 mM CaCl2] supplemented with protease inhibitors (1 mM PMSF, 2 μM pepstatin A, 10 μM leupeptin) containing 1.5% (w/v) digitonin for 30 min at 4°C. The extracts were clarified by centrifugation at 21,000 g for 30 min at 4°C, transferred into tubes containing FLAG resin (GenScript), and rotated for 2 hrs at 4°C. Post-binding, the resin was sequentially washed with lysis buffer base containing 0.1% (w/v) digitonin, high-salt wash buffer [0.1% (w/v) digitonin, 20 mM Tris-Cl (pH 7.4), 250 mM NaCl, 20 mM imidazole, 1 mM CaCl2, and 10% (v/v) glycerol], and once again with lysis buffer base containing 0.1% (w/v) digitonin. Resin-bound proteins were released by boiling in 1X reducing sample buffer and loaded onto 10 to 16% SDS-PAGE gels.

### Statistical analysis

Immunoblots were quantified using Image J and ImageLab (BIO-RAD) software. Statistical analyses were performed using Graphpad Prism (ver. 9.3). The statistical tests performed, sample sizes, and determined P values are provided in the figure or its accompanying legend.

Representative blot images from at least three independent experiments performed on at least three separate days are presented in the figures.

### Simulation methods

#### System generation

MD simulations and FEP calculations were employed to quantify the binding affinity between the Human ANT1 protein and CL. The crystal structure of the Human ANT1 protein has not been experimentally determined, therefore, a full atomic model of the Human ANT1 protein was constructed using the homology module (Prime) of the Schrödinger suite^65–67^. The sequence alignment of Human ANT1 with Bovine ANT1 and Yeast AAC2 is shown in fig. S14A. Human ANT1 is highly homologous to Bovine ANT1 (96% sequence identity), while it shares 50 – 54% sequence identify with the Yeast AAC proteins (AAC1-3). Consequently, the crystal structure of ANT1 from Bos taurus (PDB ID: 1OKC) was used as a homology modeling template and Schrödinger prime was used to build the Human model. Structural Analysis and Validation server (SAVES - https://saves.mbi.ucla.edu/) and PROCHECK^68^ was used to validate the generated homology model. Many steps of loop refinement using Schrödinger^65^ and solvent minimization (GROMACS 2020.4^69^) were performed to improve the overall quality of the structure. The final obtained high resolution (quality factor 97%) Human ANT1 protein structure is shown in the fig. S14. The quality of the obtained 3D model was assessed using the Ramachandran plot (PROCHECK) (fig. S14D). It is observed that none of the residues were lying in the disallowed regions, indicating the predicted Human ANT1 model is high quality. The Human ANT1 protein was inserted into a bilayer composed of 80% POPC (1-palmitoyl-2-oleoyl-sn-glycero-3- phosphocholine) and 20% CL (TLCL^-2^) lipids (1’-[1,2-dilinoleoyl-sn-glycero-3-phospho]-3’-[1,2- dilinoleoyl-sn-glycero-3-phospho]-glycerol) using the CHARMM-GUI membrane builder module^70, 71^. TIP3P water molecules were used to solvate the system and potassium ions were added to neutralize the total charge of the system. A sample representation of the system setup is shown in fig. S11.

#### Molecular dynamics (MD) simulation protocol

All-atom MD simulations were performed using GROMACS 2020.4^69^ with CHARMM36 force field parameters for protein and lipid molecules^72, 73^. CHARMM-GUI generated protocol was implemented to equilibrate the system. In detail, 20,000 steps of steepest descent energy minimization were performed followed by two short NVT simulations of 0.5 nanoseconds (ns) each, with a time step of 1 femtosecond (fs). Harmonic positional restraints were employed on the protein and lipid atoms during the NVT simulations; the force constants were reduced in the second NVT simulation, Subsequently, four steps of restrained NPT equilibrium simulations (0.5 ns each) were performed, where the restraints were reduced/released in each successive simulation step. The timestep was 1 fs in the first NPT simulation and 2 fs in all subsequent simulations. The Berendsen thermostat^74^ and barostat^75^ were used to maintain the system at 300 K and 1 bar pressure with a coupling constant of 1 and 5 ps, respectively. Production NPT simulations were performed for one microsecond (µs) with a time step of 2 fs. The coordinates of the trajectory were saved for every 50 picoseconds (ps). Nosé-Hoover thermostat^76^ and a Parrinello-Rahman barostat^77^ were used to maintain the temperature and pressure of the system at 300 K and 1 bar, respectively. The temperature and pressure coupling time constants were set to 1 and 5 ps, correspondingly. Semi-isotropic pressure coupling was employed with the compressibility factor of 4.5 × 10−5 bar^-1^. Long-range electrostatics were calculated with the Particle-Mesh Ewald (PME) method^78, 79^ and the short-range van der Waals and electrostatic interactions were calculated with a cut-off distance of 12 Å and used a switching distance of 10 Å. The LINCS algorithm^80^ was used to constrain the hydrogen atoms involved in covalent bonds.

To understand the effects of ANT1 protein mutations on the binding of CL, two human ANT1 mutant systems were studied in which the lysine 141 residue was mutated to phenylalanine (L141F) or glutamic acid (L141E). The final protein configuration from the 1 µs wild-type (WT) Human ANT1 simulation was used to generate the L141F and L141E systems using CHARMM-GUI. The mutant proteins were then inserted into a POPC:TLCL^-2^ (80:20) bilayer using CHARMM-GUI and the same MD protocol was used to equilibrate and simulate the mutant systems as the WT protein. All the performed simulations are detailed in Table S4. The figures were generated using VMD^81^ and PyMOL^82^. It was noticed that some CL lipid molecules occupied the known binding sites of AAC/ANT1 proteins^45–48^ in the equilibrated systems (prior to production MD). Therefore, we also ran a set of equilibrium simulations in which the prebound CL lipid molecules were removed from pocket 2 and simulated for 1 µs using the above-detailed MD protocol for both the WT and mutant proteins. Therefore, we describe a set of simulations in which CL is initially bound at pocket 2 (termed “prebound”) and a set of simulations in which the CL initially bound at pocket 2 has been removed (termed “unbound”).

#### Free Energy Perturbation (FEP) calculations

A representative frame (t≈0.965 µs) with CL^-2^ within the pocket 2 binding site, was selected from the WT prebound simulation as the input for the FEP calculations. Similarly, the corresponding equilibrium protein structure (1 µs) was used to generate the mutant FEP systems, i.e., L141F and L141E. For these calculations, the bound CL^-2^ lipid in pocket 2 was renamed to LIG (for ligand) and the same ligand conformation was used as a starting structure for all the simulated systems. The target lipid (LIG) was decoupled from the simulation box in the presence (“protein-ligand-membrane complex”) and or the absence of the Human ANT1 protein (“ligand-membrane complex”). To maintain charge neutrality during the decoupling of LIG, which carried a -2 charge, a Ca^+2^ ion was also decoupled from the system, so that both CL^-2^ and Ca^+2^ are being decoupled simultaneously. The free energies were estimated using the alchemical thermodynamic cycle described in figure S13. All FEP simulations were run in GROMACS 2020.4^66^, and a linear alchemical pathway was employed to decouple (λ=0 to λ=1) the LIG Lennard-Jones and coulombic interactions sequentailly^83^. The Lennard-Jones interactions were transformed with Δλ = 0.05 and the charges were annihilated through coulombic transformations with Δλ = 0.1. The LIG molecule was restrained, to maintain its relative position and orientation with respect to the ANT1 protein,^46, 83^ through two distances, two angles, and three dihedral harmonic potentials. Similar to Aldeghi et al. calculations, the above mentioned restraints were transformed using 12 non-uniformly distributed λ values for the protein-ligand-membrane complex systems^83^. For the ligand-membrane system, the free energy contribution of the restraints to the binding free energy was estimated analytically to be 8.311 kcal/mol.^46, 83, 84^ A total of 42 windows for the protein-ligand-membrane complex simulations and 31 windows for the ligand-membrane complex simulations were performed. For each window, 20,000 steps of steepest descent energy minimization were performed. Subsequently, all the windows were simulated for 0.5 ns under NVT ensemble conditions. The positional restraints were employed with a force constant of 500 and 100 kJ mol^-1^ nm^-2^ on the protein and lipid heavy atoms, correspondingly. Following NVT, a 1 ns position restrained NPT simulation was performed using Berendsen barostat^75^. A position restraint force of 50 kJ mol^-1^ nm^-2^ was applied on the protein and lipid heavy atoms and the pressure was set to 1 bar. Further, 0.5 ns of unrestrained NPT simulation was performed and the Parrinello-Rahman barostat^77^ was used to maintain the total pressure of the system. Finally, each window was simulated under NPT ensemble conditions for a production run of 15 ns. A semi-isotropic pressure coupling was performed with a coupling constant of 5 ps using the Parrinello–Rahman barostat. Langevin dynamics integrator was employed with a time-step of 2 fs. The PME algorithm was used with a Fourier grid spacing of 1.2 Å to calculate the long-range electrostatic interactions^78, 79^. The short-range non-bonded interactions were calculated with a cut-off distance of 12 Å and a switching distance of 10 Å. A PME spline order of 6 was employed with a relative tolerance set to 10^-6^. The soft-core potential (sigma = 0.3) was used to transform the Lennard Jones interactions^46, 83^. The GROMACS implemented long-range dispersion correction for energy and pressure was applied^83^. Lincs algorithm was used to constrain the H-bonds^80^. The Alchemical Analysis package^85, 86^ was used to estimate the free energy from the individual windows. The final 10 ns of simulation data was used to estimate the free energy values. Multistate Bennet acceptance ratio (MBAR) was used in the pymbar package to calculate free energies^86^. The entire FEP calculations were repeated four times for each simulated system, i.e., Human ANT1 WT, L141F, and L141E systems. The final binding free energies were estimated using Equation 1. Data from three repeats were averaged, and the standard errors were reported^83^. Wilcoxon rank sum tests were performed to asses statistical significance of the binding free energies between WT and mutants.

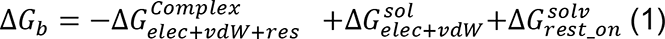

## Acknowledgements

We thank Carla Koehler (UCLA) and Jean-Paul Lasserre (University of Bordeaux) for the generous gifts of antibodies. This work was supported in part by NIH grants (R01HL108882 and R01HL165729 to S.M.C., R35GM119762 to E.R.M., Biochemistry, Cellular, and Molecular Biology Program Training Grant T32GM007445 for M.G.B. and MSTP Training Grant T32GM136577 for J.A.S.), pre-doctoral fellowships from the American Heart Association (10PRE3280013 to M.G.B. and 15PRE24480066 to O.B.O.), a post-doctoral fellowship from the Uehara Memorial Foundation (to N.S.), a post-doctoral fellowship from the American Heart Association and the Barth Syndrome Foundation (Award ID: 828058 to N.S.), and Royal Society Newton international fellowship (NIF\R1\181108 to B.S. and NIF\R1\192285 to D.K.C.)

## Author contributions

N.S. and S.M.C designed the research. N.S., D.K.C., M.G.B., B.S., O.B.O., J.A.S., T.M., Y.V., K.W., and S.M.C performed experiments. D.K.C., B.S., D.C., and C.V.R. designed native MS experiments. V.K.G., E.R.M., and N.A. designed MD simulations and V.K.G. and E.R.M. performed the experiments. N.S. wrote the manuscript draft. D.K.C., M.G.B., B.S., J.S., D.C., V.K.G., E.R.M., N.A., and S.M.C edited the manuscript.

## Declaration of interests

The authors declare no competing interests.

## Notes

### Competing Interest Statement

The authors have declared no competing interest.

