## Supplemental figures for "Conserved cardiolipin-mitochondrial ADP/ATP carrier interactions assume distinct structural and functional roles that are clinically relevant"

fig. S1

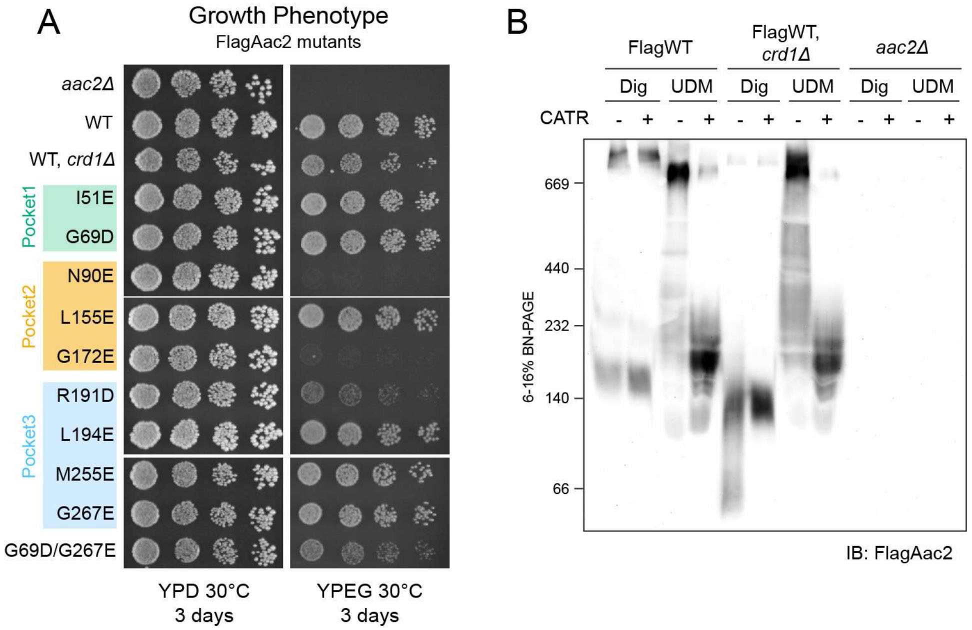


**fig. S1: Characterization of Flag-tagged WT and mutant Aac2**. (A) Growth phenotype of Flag-tagged Aac2 CL-binding mutants. Serial dilutions of indicated cells were spotted onto fermentable (YPD) and respiratory (YPEG) media and incubated at 30ºC for 3 days (n=3). (B) Mitochondria from indicated strains were mock- or pre-treated with 40 μM CATR. The treated mitochondria were then solubilized with 1.5% (w/v) digitonin or 2% (w/v) UDM, resolved by 6 to 16% blue native-PAGE and immunoblotted for Flag. Representative image from the replicates (n=3) is shown.

fig. S2

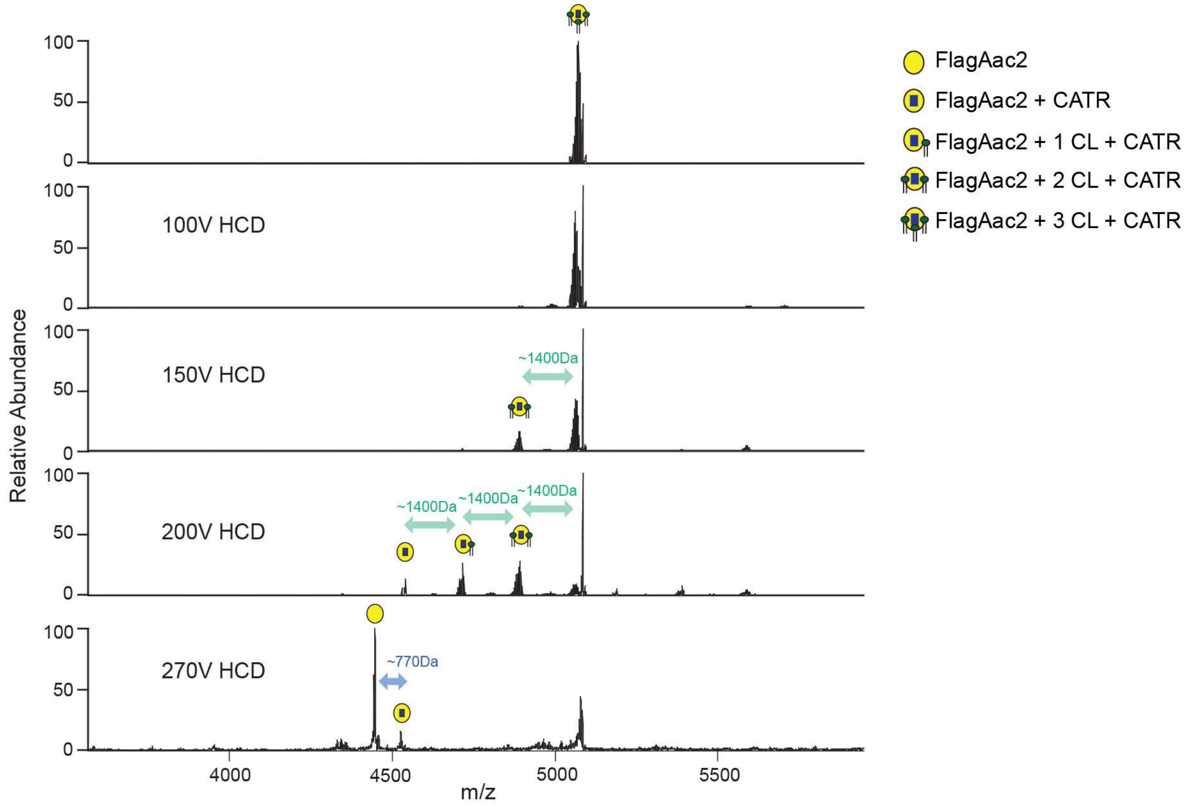


**fig. S2: Three CL molecules associated with Aac2.** Related to Fig. 2, MSMS performed against Aac2 + 3CL + CATR complex (*m/z* 5069 Da). Increased high collision dissociation (HCD) yielded spectra corresponding to CL (~1400 Da) and CATR (~770 Da).

fig. S3

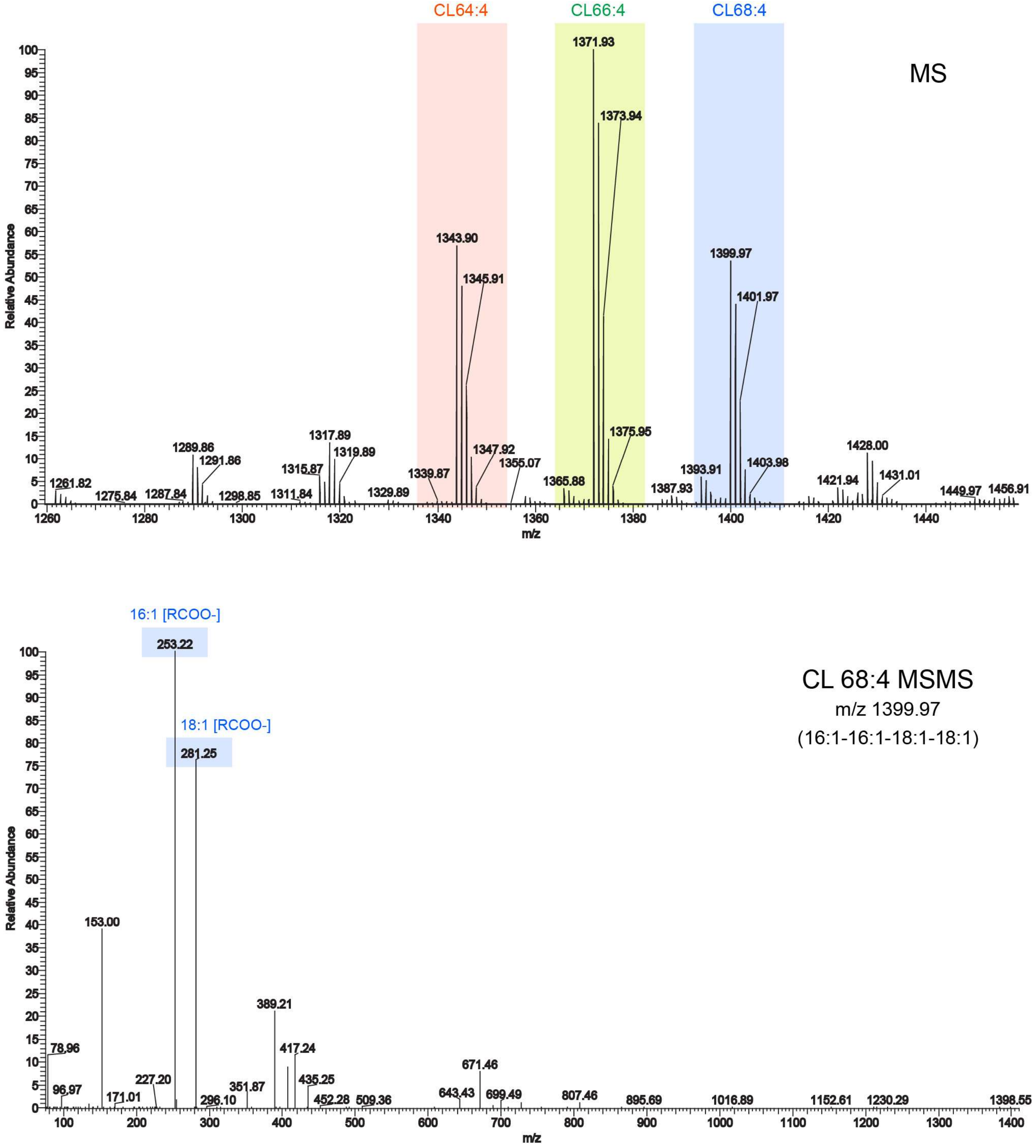


**fig. S3: CL species interacting with yeast Aac2.** Mass spectrometry (MS) analysis detected three types of CL species that co-purified with FlagAac2 from WT mitochondria (Top). MSMS performed against CL 68:4 yielded unique fragments corresponding to acyl-chains derived from CL (Bottom).

fig. S4

A

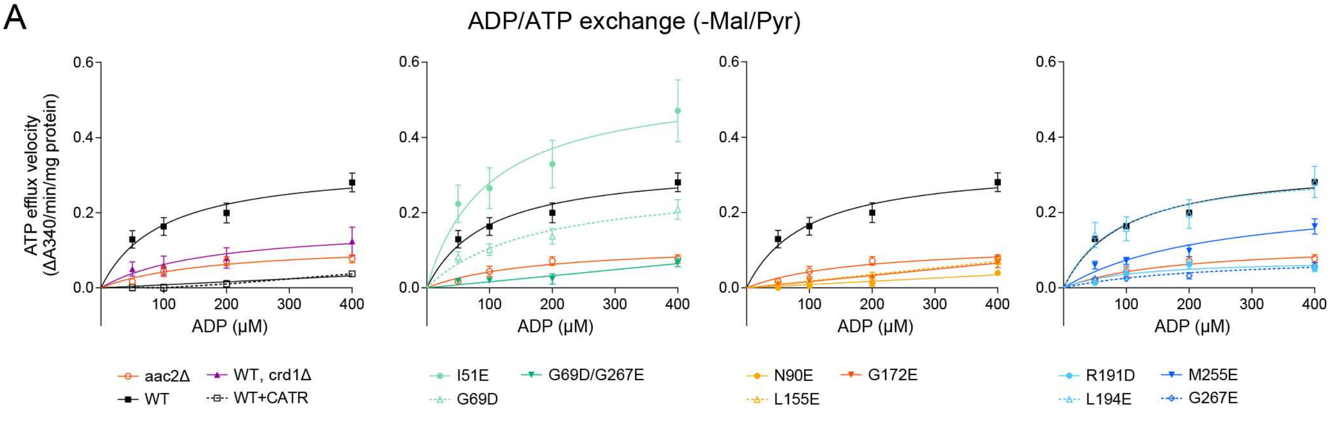


B

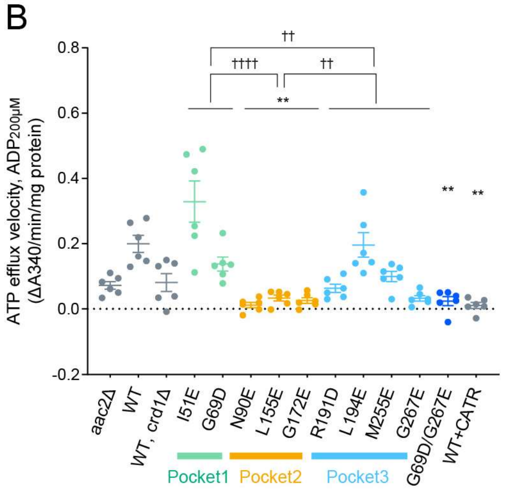


C

| | -Mal/Pyr | | | |
|---|---|---|---|---|
| | Vmax [95% CI] (μM/min/mg protein) | | Km [95%CI] (μM) | |
| aac2Δ | 0.1189 | [0.07293–0.3012] | 180.0 | [50.25–894.0] |
| WT | 0.3298 | [0.2442–0.4902] | 96.61 | [34.10–252.2] |
| WT, crd1Δ | 0.1682 | [n.d.] | 170.4 | [n.d.] |
| I51E | 0.5506 | [0.3491–1.163] | 97.38 | [14.52–485.5] |
| G69D | 0.2848 | [0.1881–0.5864] | 171.1 | [54.89–640.9] |
| N90E | n.d. | [n.d.] | n.d. | [n.d.] |
| L155E | 2.548 | [n.d.] | 14171 | [n.d.] |
| G172E | n.d. | [n.d.] | n.d. | [n.d.] |
| R191D | 0.07614 | [0.04341–0.1947] | 111.0 | [14.06–707.4] |
| L194E | 0.3212 | [0.2071–0.6405] | 89.75 | [12.82–421.9] |
| M255E | 0.2499 | [0.1548–0.7122] | 240.0 | [75.71–1231] |
| G267E | 0.08628 | [n.d.] | 234.7 | [n.d.] |
| G69D/G267E | n.d. | [n.d.] | n.d. | [n.d.] |
| WT +CATR | n.d. | [n.d.] | n.d. | [n.d.] |

**fig. S4: ADP/ATP exchange of Aac2 CL-binding mutants without respiratory substrates.** The efflux of matrix ATP was detected with isolated mitochondria as in Fig. 4A-C. The measurement was performed in the absence of malate and pyruvate (-Mal/Pyr). WT + CATR: WT mitochondria were treated with 5 μM CATR prior to the efflux reaction (n=6). (A) The linear part of the initial velocity for the ATP efflux was plotted and curve fitting performed by nonlinear regression (mean with SEM). Plots of *aac2*Δ and WT are repeated in all panels. (B) The initial linear velocity following the addition of 200 μM ADP shown as scatter plots (mean with SEM). (C) Fitted Km and Vmax values were obtained using the Michaelis-Menten equation (mean).

fig. S5

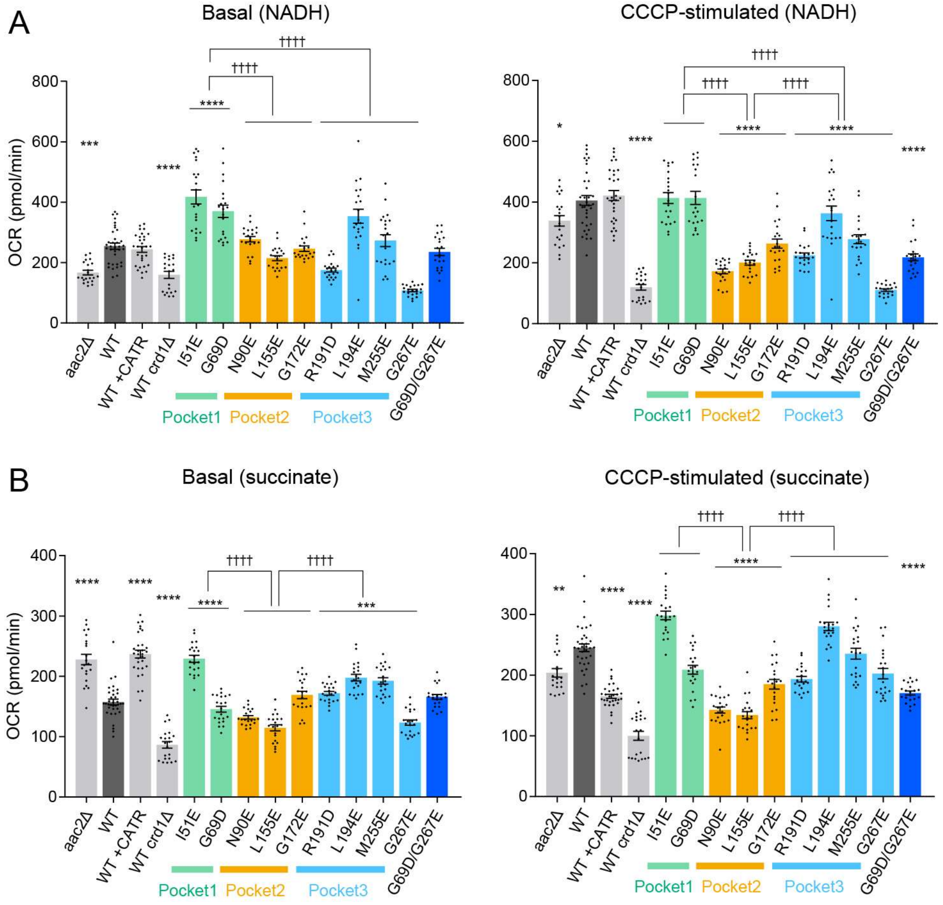


**fig. S5: Mitochondrial respiration of Aac2 CL-binding mutants.** Related to Fig. 4D-F, basal and CCCP-stimulated respirations of WT and mutant mitochondria in the presence of NADH (A) and succinate (B) were plotted as oxygen consumption rate (OCR). Mean with SEM, n=21-35. Significant differences obtained by two-way ANOVA followed by Tukey's multiple comparisons test are shown as * for comparison with WT and † for comparison between pockets; *$p<0.05$, **$p<0.01$, ***$p<0.001$, ****$p<0.0001$.

fig. S6

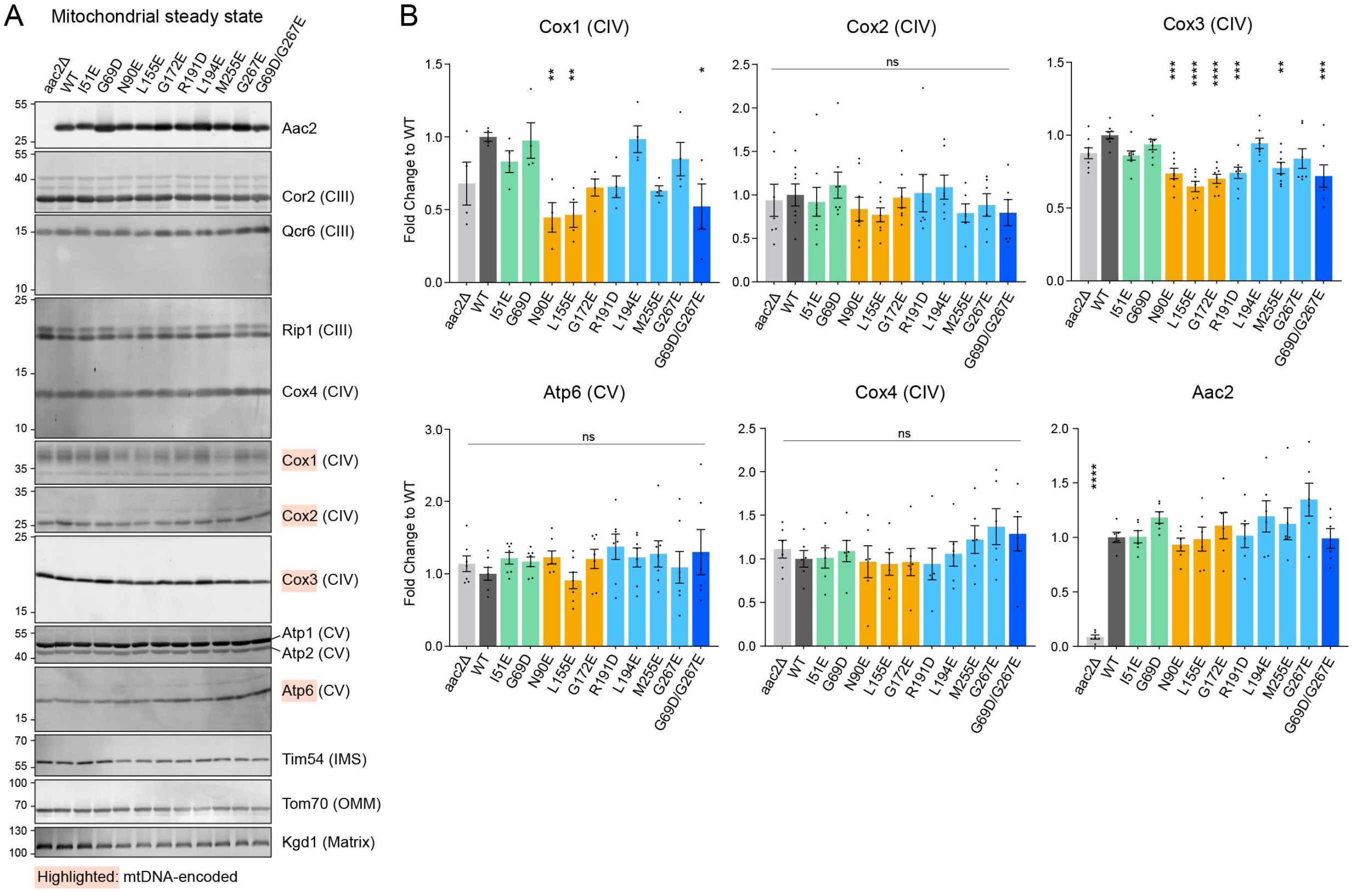


**fig. S6: The expression of respiratory complex subunits encoded in mitochondrial DNA is attenuated in Aac2 CL-binding mutants.** (A) Mitochondrial extracts were resolved by SDS-PAGE and immunoblotted for indicated proteins, including subunits of respiratory complexes III, IV, and V. (B) The expression of indicated respiratory complex subunits was quantified. Mean with SEM. Statistical differences were analyzed by one-way ANOVA followed by Dunnett's multiple comparison test; *p<0.05, **p<0.01, ***p<0.001, ****p<0.0001 (vs. WT). Representative images from the replicates (n=4-8) are shown.

fig. S7

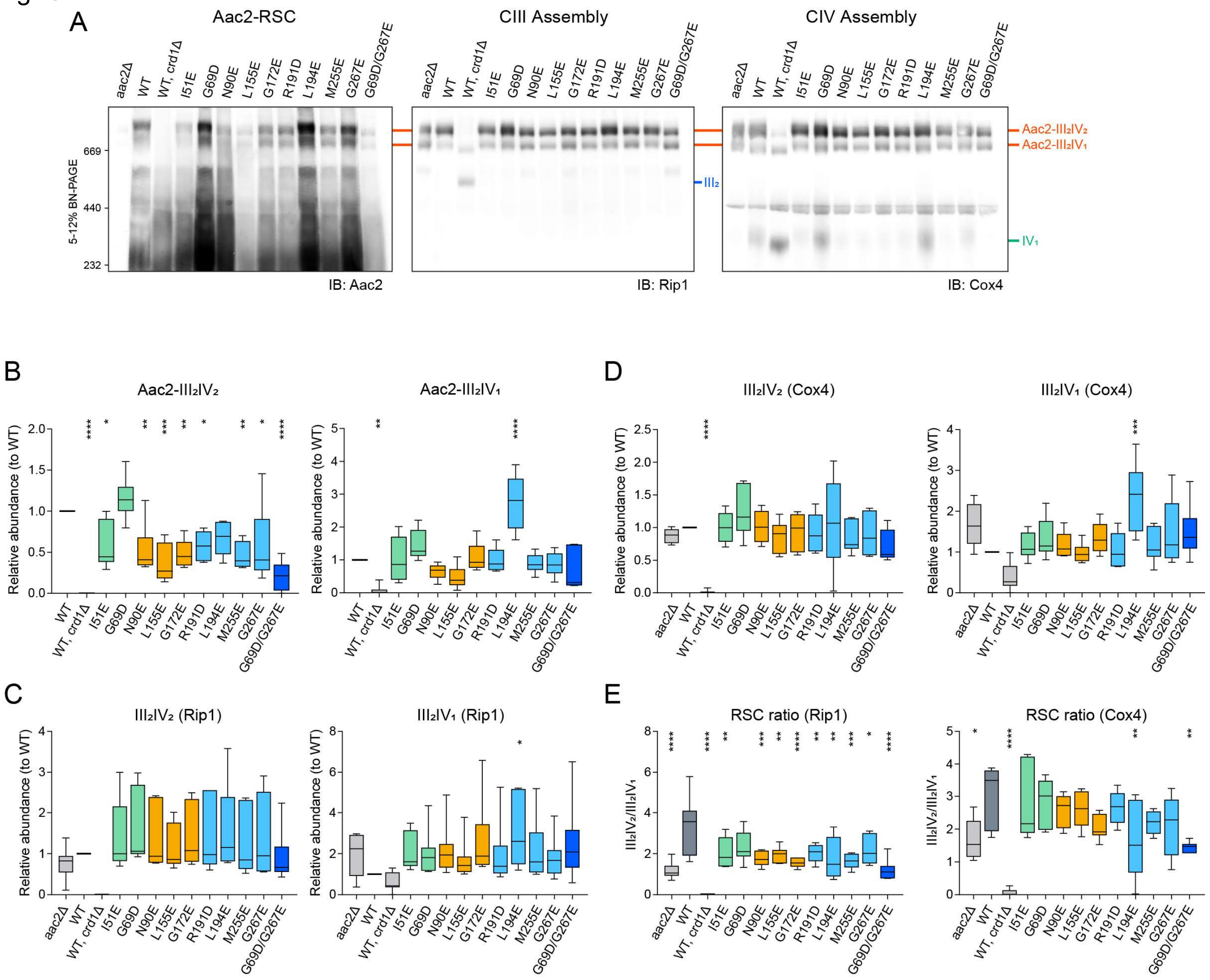


**fig. S7: Assembly of Aac2 CL-binding mutants and respiratory supercomplexes is modestly altered.** (A) WT and mutant mitochondria were solubilized with 1.5% (w/v) digitonin, resolved by 5 to 12% blue native-PAGE, and immunoblotted (IB) as indicated. RSC, respiratory supercomplex. (B-D) Quantification of assembled Aac2 (B), Rip1 (C), and Cox4 (D) within respiratory supercomplexes. (E) Ratios of respiratory supercomplexes $III_2IV_2$ and $III_2IV_1$ when detected by Rip1 and Cox4, respectively. Data are shown as box-whisker plots with the box extended from 25th to 75th percentiles and the whiskers indicating the min to max range. One-way ANOVA followed by Dunnett's multiple comparison test determined the significance; *$p<0.05$, **$p<0.01$, ***$p<0.001$, ****$p<0.0001$. Representative images from the replicates (n=5-6) are shown; images have been cropped to exclude the abundant Aac2 monomer to facilitate visualization of the Aac2-RSC complexes.

fig. S8

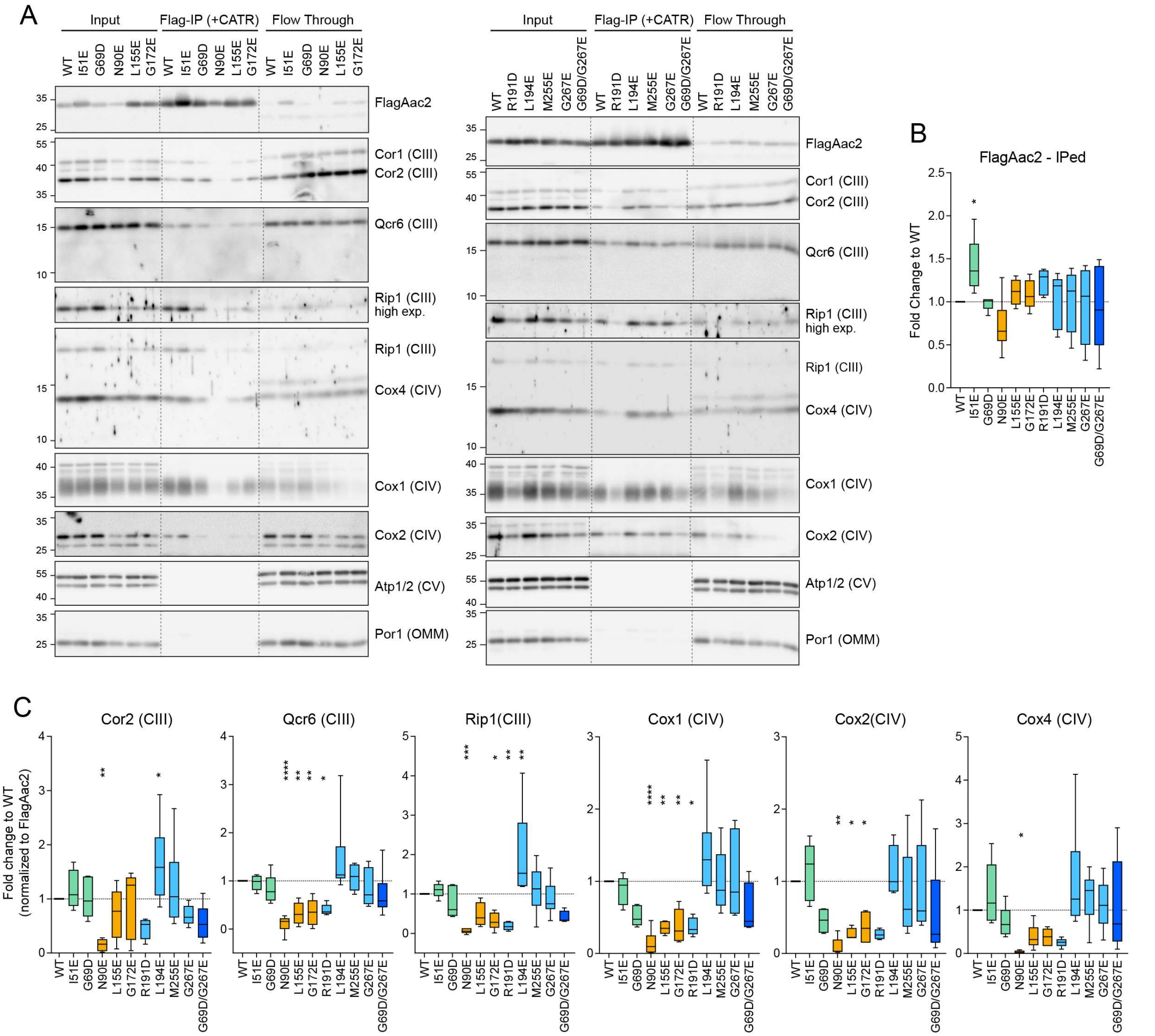


**fig. S8: Protein-protein interaction between Aac2 and respiratory complex subunits are diminished in Aac2 CL-binding mutants.** (A) Isolated mitochondria from Flag-tagged WT and mutant Aac2 strains were pre-incubated with 40 μM CATR and then solubilized with 1.5% (w/v) digitonin. The mitochondrial extracts were immunoprecipitated (IP) using anti-Flag resin. Co-purified subunits of complexes III and IV were determined by immunoblotting; Atp1/2 and Por1 served as controls. Four percent of input (intact mitochondria) and flow through (unbound) was analyzed. (B) The abundance of FlagAac2 eluted upon IP. (C) The abundance of subunits of complexes III and IV co-purified with FlagAac2 was quantified and normalized. Data are shown as box-whisker plots with the box extended from 25th to 75th percentiles and the whiskers indicating the min to max range. Statistical differences were analyzed by one-way ANOVA followed by Dunnett's multiple comparison test; *$p<0.05$, **$p<0.01$, ***$p<0.001$, ****$p<0.0001$ (vs. WT). Representative images from the replicates (n=4-12) are shown.

fig. S9

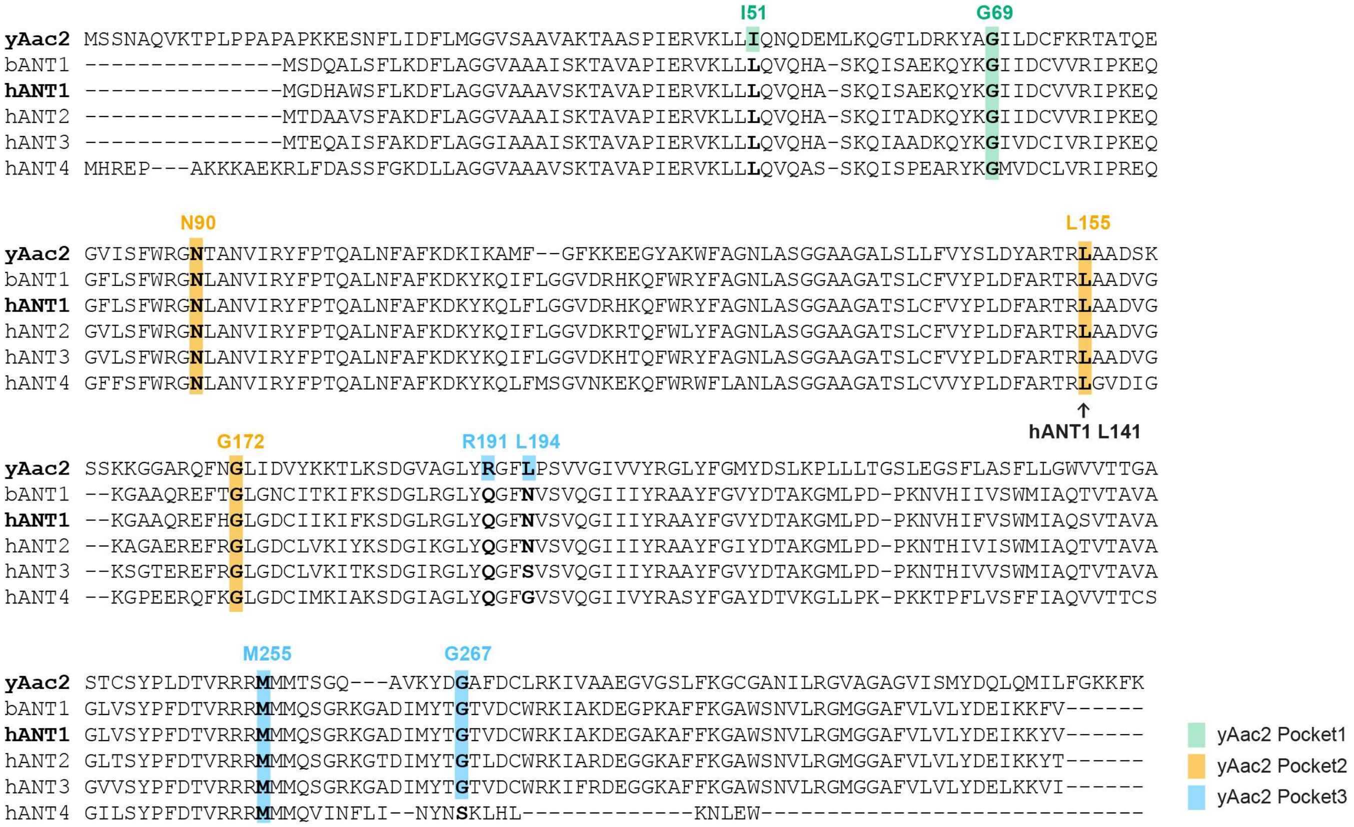


**fig. S9: CL-binding sites are conserved across species.** Amino acid sequence alignment of yeast Aac2, bovine ANT1, and human ANT isoforms. The residues designed for the Aac2 CL-binding mutants are highlighted as indicated.

fig. S10

A

```
                                         R106H                                   148-AGA/SGT-150    Y165T K171R
P12235|ANT1 ------------MGDHAWSFLKDFLAGGVAAAVSKTAVAPIERVKLLLQVQHASKQISAEKQYKGIIDCVVRIPKEQGFLSFWRGNLANVIRYFPTQALNF 89
P05141|ANT2 ------------MTDAAVSFAKDFLAGGVAAAISKTAVAPIERVKLLLQVQHASKQITADKQYKGIIDCVVRIPKEQGVLSFWRGNLANVIRYFPTQALNF 89
P12236|ANT3 ------------MTEQAISFAKDFLAGGIAAAISKTAVAPIERVKLLLQVQHASKQIAADKQYKGIVDCIVRIPKEQGVLSFWRGNLANVIRYFPTQALNF 89
Q9H0C2|ANT4 MHREPAKKKAEKRLFDASSFGKDLLAGGVAAAVSKTAVAPIERVKLLLQVQASSKQISPEARYKGMVDCLVRIPREQGFFSFWRGNLANVIRYFPTQALNF 101
                            * ** **:****:***:******************* :****: : :***::**:****:***.:*********************

P12235|ANT1 AFKDKYKQLFLGGVDRHKQFWRYFAGNLASGGAAGATSLCFVYPLDFARTRLAADVGKGAAQREFHGLGDCIIKIFKSDGLRGLYQGFNVSVQGIIIYRAA 190
P05141|ANT2 AFKDKYKQIFLGGVDKRTQFWLYFAGNLASGGAAGATSLCFVYPLDFARTRLAADVGKAGAEREFRGLGDCLVKIYKSDGIKGLYQGFNVSVQGIIIYRAA 190
P12236|ANT3 AFKDKYKQIFLGGVDKHTQFWRYFAGNLASGGAAGATSLCFVYPLDFARTRLAADVGKSGTEREFRGLGDCLVKITKSDGIRGLYQGFSVSVQGIIIYRAA 190
Q9H0C2|ANT4 AFKDKYKQLFMSGVNKEKQFWRWFLANLASGGAAGATSLCVVYPLDFARTRLGVDIGKGPEERQFKGLGDCIMKIAKSDGIAGLYQGFGVSVQGIIVYRAS 202
            ********:*:.**::..*** :* .**************.***********..*:**.  :*:*:*****::** ****: ******.*******:***:

                                      T227V                T247A           A262F
P12235|ANT1 YFGVYDTAKGMLPDPKNVHIFVSWMIAQSVTAVAGLVSYPFDTVRRRMMMQSGRKGADIMYTGTVDCWRKIAKDEGAKAFFKGAWSNVLRGMGGAFVLVLY 291
P05141|ANT2 YFGIYDTAKGMLPDPKNTHIVISWMIAQTVTAVAGLTSYPFDTVRRRMMMQSGRKGTDIMYTGTLDCWRKIARDEGGKAFFKGAWSNVLRGMGGAFVLVLY 291
P12236|ANT3 YFGVYDTAKGMLPDPKNTHIVVSWMIAQTVTAVAGVVSYPFDTVRRRMMMQSGRKGADIMYTGTVDCWRKIFRDEGGKAFFKGAWSNVLRGMGGAFVLVLY 291
Q9H0C2|ANT4 YFGAYDTVKGLLPKPKKTPFLVSFFIAQVVTTCSGILSYPFDTVRRRMMMQSGEA--KRQYKGTLDCFVKIYQHEGISSFFRGAFSNVLRGTGGALVLVLY 301
            *** ***.**:**.**:. :.:*::*** **: :*: ****************.   .  *.**:**: ** :.** .:**:**:****** ***:*****

P12235|ANT1 DEIKKYV------- 298
P05141|ANT2 DEIKKYT------- 298
P12236|ANT3 DELKKVI------- 298
Q9H0C2|ANT4 DKIKEFFHIDIGGR 315
            *::*:
```

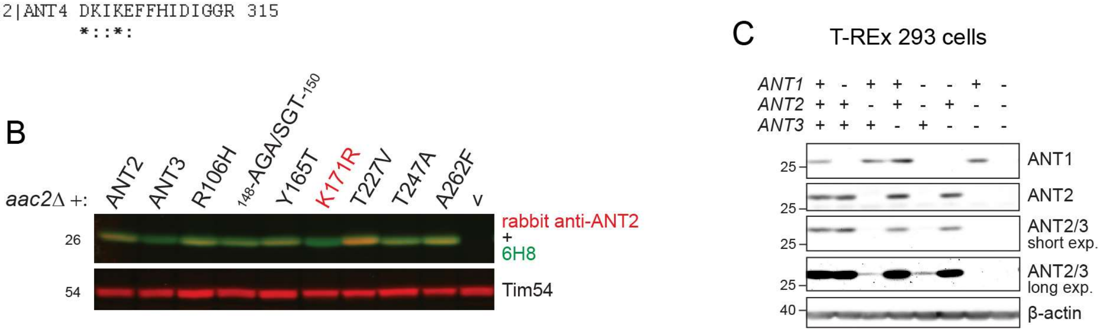


**fig. S10: Endogenous expression of three ANT isoforms was absent in ant$^{null}$ cells.** (A, B) Peptide mapping of ANT2 antisera. (C) The expression of three ANT isoforms was detected in whole cell extracts by immunoblot (n=5). The absence of ANT1, ANT2, and ANT3 was confirmed in ant$^{null}$ cells (right-most lane). Representative images from the indicated replicates in B and C are shown.

fig. S11

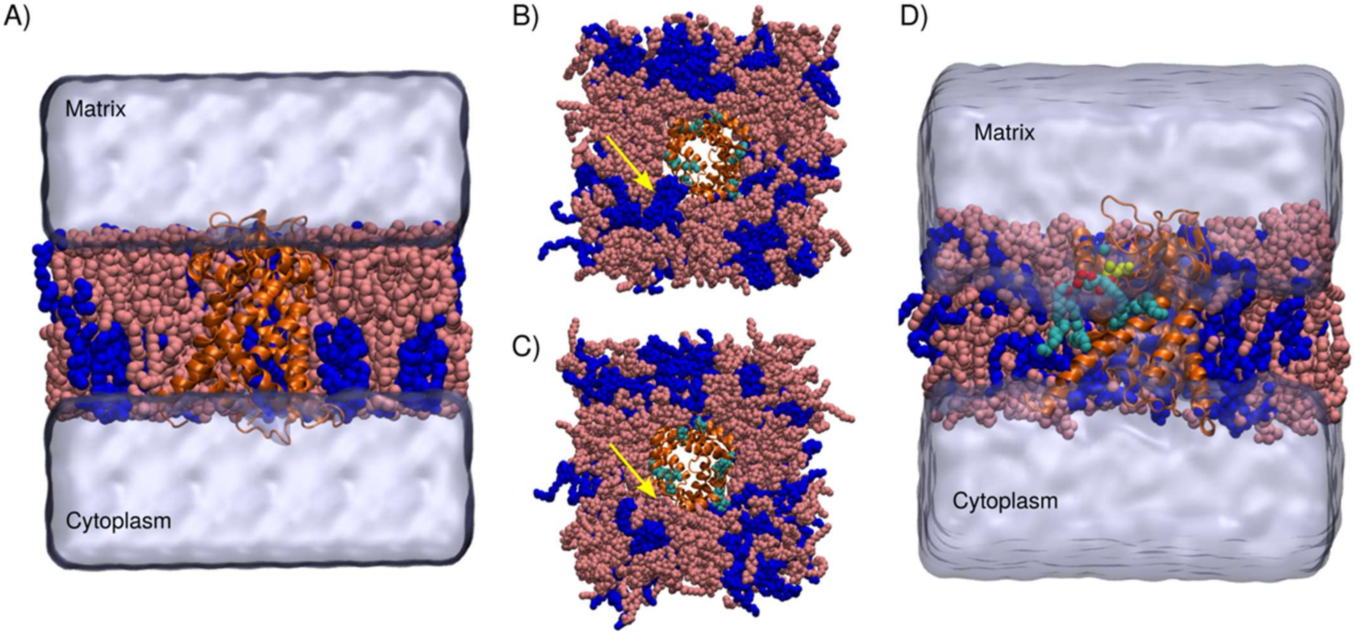


**fig. S11: Human ANT1 simulation system setup (c-state).** (A) The ANT1 protein (Orange cartoon), POPC (pink), and TLCL2 (tetralinoleoyl-cardiolipin $(18{:}2)_4$ in di-anionic form) (blue) were solvated in water (iso-blue surface). The top view (matrix view) of the ANT1 system setup for the equilibrium prebound (B) and equilibrium unbound (C) simulations. Yellow arrows point to the presence or absence of CL lipid, around pocket 2. (D) Human ANT1 free energy perturbation (FEP) calculation system setup showing the "ligand CL", LIG (head group oxygen atoms in red, phosphorous atoms in green and acyl chain atoms in cyan van der Waals representation). The front portion of the membrane, hydrogen atoms, and the water molecules were removed for clarity.

fig. S12

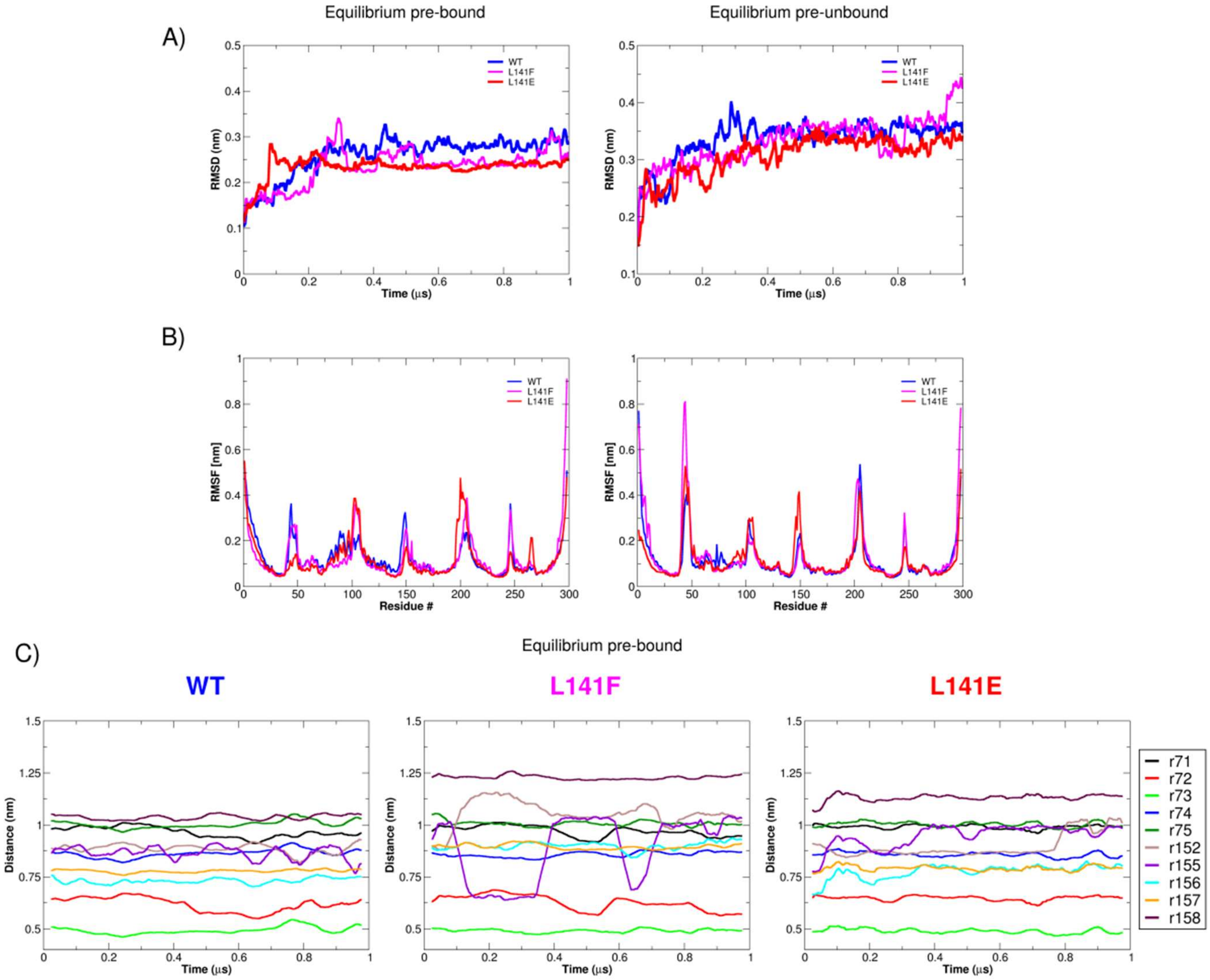


**fig. S12: ANT1 protein dynamics during MD simulations.** (A) Root-mean-squared deviation (RMSD) in prebound (left) and unbound (right) 1 µs simulations; 100 frame running averaging was performed to smooth the curves. (B) Root-mean-squared fluctuations (RMSF) for prebound (left) and unbound (right) simulations. (C) Calculated distances between the Cα atoms of residue 141 with that of the selected neighboring and pocket 2 binding site residues (residues 71, 72, 73, 74, 75, 152, 155, 156, 157, and 158) during prebound simulations.

fig. S13

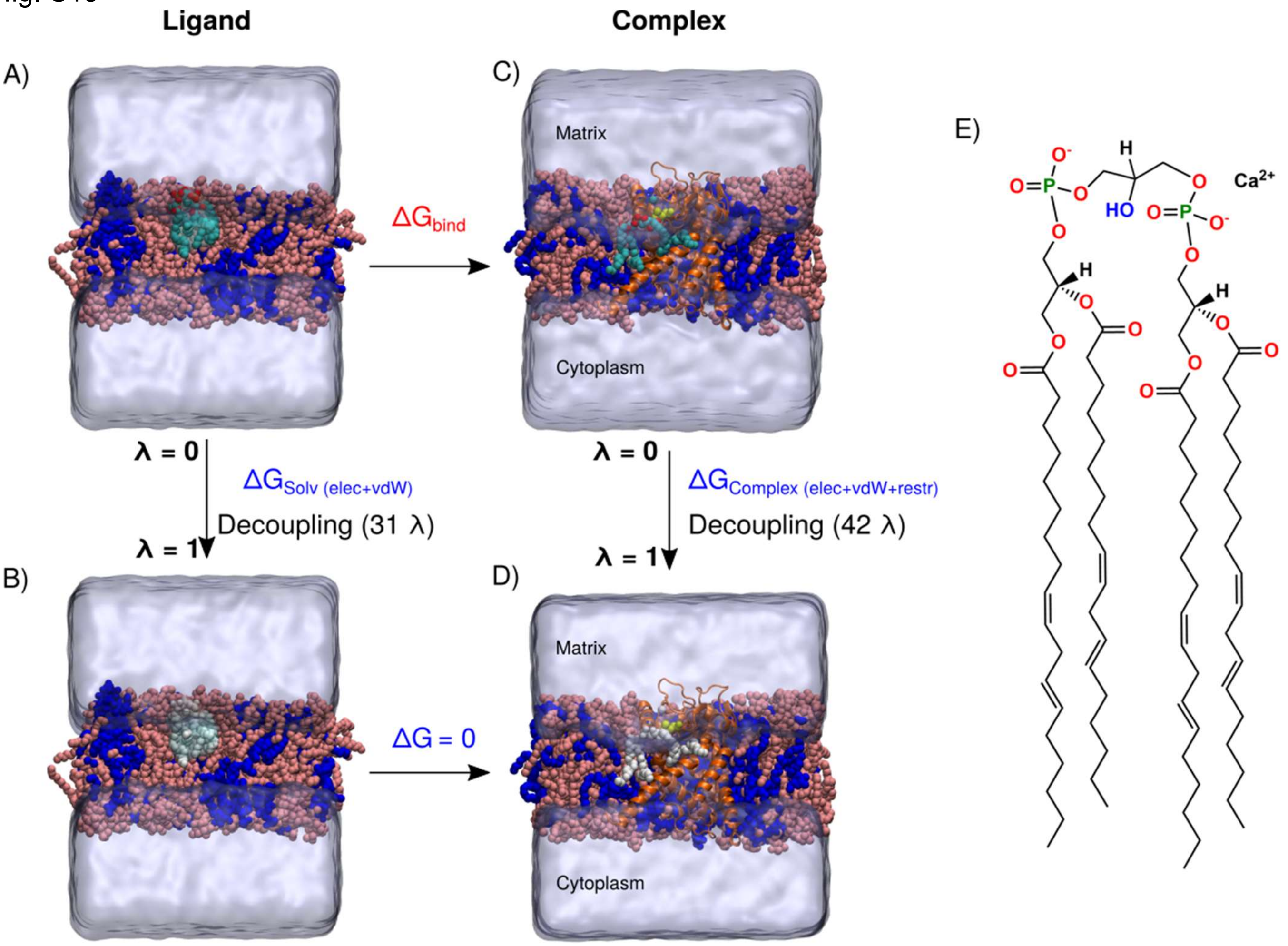


**fig. S13: FEP thermodynamic cycle.** (A) The fully integrated CL LIG in a bilayer environment is transformed into a completely non-interacting ligand (B, white) during a series of 31 equilibrium simulations in which corresponding electrostatic and van der Waals interactions are scaled to zero. The fully interacting LIG at the top right (C) is transformed into a completely non-interacting ligand (D, white) in the presence of ANT1 membrane protein during a series of 42 equilibrium simulations in which corresponding restraints, electrostatic, and van der Waals interactions are scaled to zero. The ANT1 protein (Orange cartoon), the POPC and TLCL2 membrane lipids (pink and blue van der Waals representation), and ligand LIG (head group oxygen atoms in red, phosphorous atoms in green and acyl chain atoms in cyan van der Waals representation) were solvated in water (iso-blue surface). (E) 2D structure of LIG used in the present study FEP calculations including the $Ca^{+2}$ which was simultaneously decoupled with LIG to maintain charge neutrality.

fig. S14

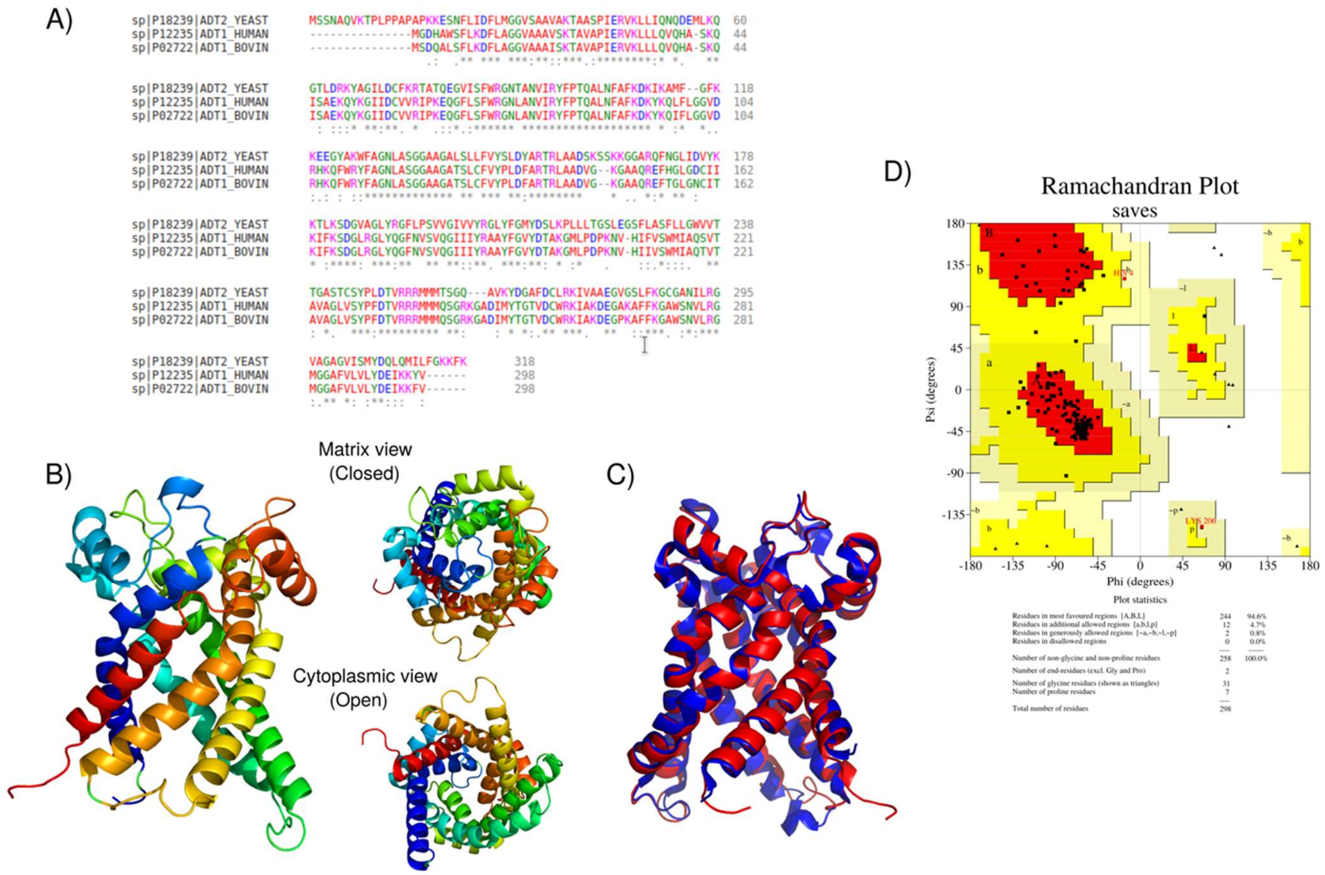


**fig. S14: Human ANT1 homology modeling.** (A) Sequence alignment of Human ANT1 with Bovine ANT1 and Yeast AAC2. Bovine ANT1 (PDB ID: 1OKC and 2C3E) was used as a template. (B) Cartoon representation of the generated Human ANT1 homology model (97% Quality factor). (C) Overlap structures of Bovine ANT1 (1OKC) and Human ANT1 model proteins. (D) Ramachandran plot of ANT1 model from Human generated by PROCHECK: in which 94.6 % residues in favorable regions; 4.7 % residues in additional allowed regions; 0.8 % residues in generously allowed regions; 0% residues in disallowed regions.

**Table S1 Primers used to generate yeast mutant constructs.**

| Target | Type | Sequence (5'-3') |
|---|---|---|
| Aac2 5' UTR (SalI) | Forward | ACGCGTCGACGAGCACTGTTTCCAATGGAG |
| Aac2 3' End (NotI) | Reverse | GTGGCGGCCGCTCTTATTTGAACTTCTTACCAAAC |
| Aac2 I51E | Forward | ACTTTTGGAACAAAACCAAGATGAAATGTTAAAAC |
| Aac2 I51E | Reverse | ATCTTGGTTTTGTTCCAAAAGTTTAACTCTTTCGATGG |
| Aac2 G69D | Forward | AAAATACGCAGATATCTTAGACTGTTTCAAGAGAAC |
| Aac2 G69D | Reverse | CAGTCTAAGATATCTGCGTATTTTCTGTCCAAAG |
| Aac2 N90E | Forward | GGAGAGGTGAGACTGCTAACGTTATCCGTTATTTC |
| Aac2 N90E | Reverse | GTTAGCAGTCTCACCTCTCCAGAATGAGATAAC |
| Aac2 L155E | Forward | CAAGAACTAGAGAAGCTGCTGACTCCAAGTC |
| Aac2 L155E | Reverse | AGCAGCTTCTCTAGTTCTTGCATAATCCAAAG |
| Aac2 G172E | Forward | GTCAATTCAACGAATTGATCGATGTCTACAAGAAG |
| Aac2 G172E | Reverse | CGATCAATTCGTTGAATTGACGAGCACC |
| Aac2 R191D | Forward | GGTCTTTACGACGGTTTCTTACCTTCTGTCGTTG |
| Aac2 R191D | Reverse | AAGAAACCGTCGTAAAGACCAGCAACACCATC |
| Aac2 L194E | Forward | CAGAGGTTTCGAACCTTCTGTCGTTGGTATTG |
| Aac2 L194E | Reverse | CAGAAGGTTCGAAACCTCTGTAAAGACCAG |
| Aac2 M255E | Forward | AAGAAGAGAGATGATGACCTCCGGTCAAGC |
| Aac2 M255E | Reverse | GAGGTCATCATCTCTCTTCTTCTAACGGTATCCAATG |
| Aac2 G267E | Forward | GTTAAGTACGACGAAGCCTTTGACTG |
| Aac2 G267E | Reverse | AAAGGCTTCGTCGTACTTAACAGC |
| Aac2 L155F | Forward | AAGAACTAGATTCGCTGCTGACTCCAAGTCCTC |
| Aac2 L155F | Reverse | GAGTCAGCAGCGAATCTAGTTCTTGCATAATCC |

| | | |
|---|---|---|
| Aac2 N-term Flag | Forward | ATGGATTATAAAGATGATGACGATAAAATGTCTTCCAACGCCCAAGTC |
| Aac2 N-term Flag | Reverse | TTTATCGTCATCATCTTTATAATCCATGGCTATTTGCTTATATGTATGTTAATGT |

**Table S2 Primers used to generate human mutant constructs.**

| Target | Type | Sequence (5'-3') |
|---|---|---|
| ANT1 5' Flag (HindIII) | Forward | CCCAAGCTTATGGATTATAAAGATGATGACGATAAAATGGGTGATCACGCTTGGAG |
| ANT1 3' End (NotI) | Reverse | ATTTGCGGCCGCTTAGACATATTTTTTGATCTC |
| ANT1 L141E | Forward | GCTAGGACCAGGGAGGCTGCTGATGTGGGCAAG |
| ANT1 L141E | Reverse | ATCAGCAGCCTCCCTGGTCCTAGCAAAGTCCAGC |
| ANT1 L141F | Forward | GCTAGGACCAGGTTCGCTGCTGATGTGGGCAAG |
| ANT1 L141F | Reverse | ATCAGCAGCGAACCTGGTCCTAGCAAAGTCCAGC |

**Table S3 Antibodies used in this study.**

| Antibodies | Source | Identifier | Used in |
|---|---|---|---|
| Flag, mouse monoclonal (M2) | Sigma-Aldrich | F3165 | fig. S1B |
| Flag, mouse monoclonal (12C6c) | Developmental Studies Hybridoma Bank (DSHB) | RRID:AB_2890618 | Fig. 5C, E; Fig. 6B, C; |
| Flag, rabbit polyclonal | Sigma-Aldrich | SAB4301135 | fig. S8A |
| Aac2, mouse monoclonal (6H8) | Panneels et al. 2003, Biochem Biophys Res Commun [36] | 6H8 | Fig. 1B, D; Fig. 3A, B; fig. S7A; fig. S10B |
| Tom70, rabbit polyclonal | Riezman et al. 1983, EMBO J [87] | 7305 | Fig. 1B, D, Fig. 5C; fig. S6A |
| Atp1/2, rabbit polyclonal | Maccecchini et al. 1979, Proc Natl Acad Sci [88] | UY3-T | fig. S6A; fig. S8A |
| Por1, rabbit polyclonal | Daum et al. 1982, J Biol Chem [89] | 425 | fig. S8A |
| Kgd1, rabbit polyclonal | Glick et al. 1992, Cell [90] | 453-3 | Fig. 1B, Fig. 5C; fig. S6A |
| Cor2, rabbit polyclonal | Glick et al. 1992, Cell [90] | CC2-T | fig. S6A; fig. S8A |
| Cox1, rabbit polyclonal | Dowhan et al. 1985, EMBO J [91] | DD2-4 | fig. S6A; fig. S8A |
| Cox2, rabbit polyclonal | Poyton et al. 1975, J Biol Chem [92] | 173 | fig. S6A; fig. S8A |

| | | | |
|---|---|---|---|
| Cox3, mouse monoclonal (DA5BC4) | Invitrogen | 459300 | fig. S6A |
| Cox4, rabbit polyclonal | Baile et al. 2013, Mol Biol Cell [93] | MGB65 | fig. S6A; fig. S7A; fig. S8A |
| Rip1, rabbit polyclonal | Baile et al. 2013, Mol Biol Cell [93] | MGB71 | fig. S6A; fig. S7A; fig. S8A |
| Qcr6, rabbit polyclonal | Baile et al. 2013, Mol Biol Cell [93] | MGB73 | fig. S6A; fig. S8A |
| Atp6, rabbit polyclonal | Kabala et al. 2014, Biochimie [94] | N/A | fig. S6A |
| Taz, rabbit polyclonal | Claypool et al. 2006, J Cell Biol [59] | 4248 | Fig. 1D |
| Abf2, rabbit polyclonal | Calzada et al. 2019, Nat Commun [55] | 5477 | Fig. 1D |
| Tim54, rabbit polyclonal | This study | 7303 | fig. S6A; fig. S10B |
| β-actin, mouse monoclonal | Sigma-Aldrich | A5441; RRID:AB_476744 | Fig. 6B, fig. S10C |
| GRP75, mouse monoclonal | Antibodies Incorporated | 75-127; RRID: AB_2120479 | Fig. 6B |
| ANT1, mouse monoclonal (1F3F11) | Lu et al. 2017, Mol Cell Biol [58] | N/A | fig. S10C |
| ANT2, rabbit polyclonal | Acoba et al. 2021, Cell Rep [56] | 5695 | fig. S10B, C |
| ANT2/3, mouse monoclonal (5H7) | Panneels et al. 2003, Biochem Biophys Res Commun [36] | N/A | fig. S10C |
| HRP-conjugated secondary, goat anti-rabbit IgG (H+L) | Thermo Fisher Scientific | 31460; RRID:AB_228341 | Fig. 1B, D; Fig. 3A, B; Fig. 5C, E; Fig. 6 B, C; fig. S1B; fig. S7A; fig. S8A; fig. S10B |
| HRP-conjugated secondary, goat anti-mouse IgG (H+L) | Thermo Fisher Scientific | 62-6520; RRID:AB_2533947 | Fig. 1B, D; Fig. 3A, B; Fig. 5C, E; Fig. 6 B, C; fig. S1B; fig. S7A; fig. S8A; fig. S10B |
| Daylight 650 conjugated secondary, goat anti-rabbit IgG (H+L) | Invitrogen | 84546 | fig. S6 |
| Daylight 550 conjugated secondary, goat anti-mouse IgG (H+L) | Invitrogen | 84540 | fig. S6 |

**Table S4 Overview of the simulation setup and details.**

| Simulation Methods | System | Simulation length |
|---|---|---|
| Equilibrium<br>Pocket 2 CL prebound | WT | 1 X 1 μs |
| | L141F | 1 X 1 μs |
| | L141E | 1 X 1 μs |
| Equilibrium<br>Pocket 2 CL unbound | WT | 1 X 1 μs |
| | L141F | 1 X 1 μs |
| | L141E | 1 X 1 μs |
| Free Energy Perturbations (FEP) | WT (42 X 15 ns) | 4 X 0.63 μs = 2.52 μs |
| | L141F (42 X 15 ns) | 4 X 0.63 μs = 2.52 μs |
| | L141E (42 X 15 ns) | 4 X 0.63 μs = 2.52 μs |
| | Ligand (31 X 15 ns) | 4 X 0.465 μs = 1.86 μs |
| | | Total = 15.42 μs |
